# A simple framework for agent-based modeling with extracellular matrix

**DOI:** 10.1101/2022.11.21.514608

**Authors:** John Metzcar, Ben S. Duggan, Brandon Fischer, Matthew Murphy, Randy Heiland, Paul Macklin

**Affiliations:** Intelligent Systems Engineering, Indiana University, 700 N. Woodlawn, Bloomington, 47408, Indiana, United States of America; Informatics, Indiana University, 901 E. Tenth Street, Bloomington, 47408, Indiana, United States of America; Computer Science, Indiana University, 700 N. Woodlawn, Bloomington, 47408, Indiana, United States of America

**Keywords:** extracellular matrix, agent-based modeling, collective migration, stigmergy, fibrosis, cancer and basement membrane invasion

## Abstract

Extracellular matrix (ECM) is a key component of the cellular microenvironment and critical in multiple disease and developmental processes. Representing ECM and cell-ECM interactions is a challenging multiscale problem as they span molecular-level details to tissue-level dynamics. While several computational frameworks exist for ECM modeling, they often focus on very detailed modeling of individual ECM fibers or represent only a single aspect of the ECM. Using the PhysiCell agent-based modeling platform, we developed a framework of intermediate detail with the ability to capture bidirectional cell-ECM interactions. We represent a small region of ECM, an ECM element, with three variables describing its local microstructure: anisotropy, density, and overall fiber orientation. To spatially model the ECM, we use an array of ECM elements. Cells remodel local ECM microstructure and in turn, local microstructure impacts cellular motility. We demonstrate the utility of this framework and reusability of its core cell-ECM interaction model through examples in cellular invasion, wound healing, basement membrane degradation, and leader-follower collective migration. Despite the relative simplicity of the framework, it is able to capture a broad range of cell-ECM interactions of interest to the modeling community. Furthermore, variables representing the ECM microstructure are accessible through simple programming interfaces. This allows them to impact cell behaviors, such as proliferation and death, without requiring custom code for each interaction, particularly through PhysiCell’s modeling grammar, enabling rapid modeling of a diverse range of cell-matrix biology. We make this framework available as a free and open source software package at https://github.com/PhysiCell-Models/collective-invasion.

## 1 Introduction

Extracellular matrix (ECM), the material in which cells live, plays a role in a host of biological processes, from facilitating cell-cell communication to tissue development and organization. It is diverse with composition-dependent biomechanical and biochemical properties that vary greatly across locations within the body [1]. It is a living material, actively maintained and modified by cells such as fibroblasts in wound healing and cancer cells in local cellular invasion [1–3]. Its properties are formed from multiple interacting fibers, for example, through chemical crosslinking of ECM components [2]. These fibers are composed of proteins and polysaccharides which are typically organized into several large families: proteoglycans, laminins, fibronectin, and fibrous proteins, each of which can be further grouped by physical and chemical properties [1]. Fibrous proteins include the collagens, the primary structural component of interstitial ECM, and elastin, which provides elastic properties to the ECM. Basement membrane ECM, which surrounds and separates functional units of tissue, is typically composed of collagens interwoven with laminins [2]. Additionally, the ECM includes signaling molecules and many attachment points for cell-ECM interaction [2, 3]. While cells generate and maintain ECM, ECM also impacts cellular behavior. In particular, matrix density can hinder or enhance cell migration depending on cell type [4, 5]. Additionally, ECM fiber orientations can guide cellular migration through contact guidance, in which cells follow ECM, reorienting their motion to travel parallel to ECM fibers [6–12]. For example, Carey et al. observed that fiber alignment induced anisotropic cell morphodynamics and spatial probing, guiding them along the axis of alignment [13]. Furthermore, ECM-based signaling, both through mechanosensing and cell receptor binding to ECM molecules, can impact cell cycle and death [14]. Cell differentiation can also be impacted by ECM [15]. Finally, changes in ECM microstructure are associated with disease processes, in particular aspects of cancer progression, cancer metastasis, and scarring [9, 10, 16, 17].

Given the importance of ECM in biological processes, there are multiple models and frameworks for exploring ECM and cell-matrix interactions across a range of contexts such as cancer, wound healing, and fibrosis [18–21]. To provide a brief overview of previous works, we place these efforts in four categories: discrete force-based, continuous force-based, ECM density, and multicomponent that have both ECM density and orientation.

Examples of the discrete force-based frameworks include Schlüter et al. and Nöel et al. where migrating cells and individual matrix fibers are embodied as agents with direct forces dictating motility and realignment respectively [22, 23]. Macnamara et al. also adopt this paradigm and couple it with a model of blood vessel-cell interactions [24]. Other representations calculate forces over a network of individual spring-like elements. Zhu et al. construct ECM from a hook-and-spring network and Tozluoğlu et al. model individual fiber filaments as hook-and-spring networks [25, 26]. Tsingos et al. model a network of ECM fibers as a bead-spring chain model and couple this with a cellular-Potts agent-based model [27, 28].

Continuous force-based models represent ECM as a continuously deformable material. Examples include van Oers et al., who use a finite element model of ECM coupled with a cellular-Potts model of cells and look at how stiffness and deformability of the ECM impact cell migration [29]. Ahmadzadeh et al. use a stress-strain relationship to deform fibers and incorporate cell adhesions to investigate cellular level motility and morphology [30].

ECM density approaches often use partial-differential equations (PDEs) to model ECM density. Anderson and Chaplain use a continuous method that leads to a hybrid discrete-continuous model to study the interplay between angiogenesis and ECM [31]. Zeng et al. use a fibronectin concentration in developmental biology applications [32]. Daube et al. use a PDE for a scalar ECM density field in the context of sprouting angiogenesis in tumors [33]. Trucu et al. model invasion as a moving boundary problem with ECM represented as a scalar field [34]. Building off that work, Shuttleworth et al. add non-local effects as well as ECM orientation and degradation to the invasive boundary [35, 36]. Gonçalves and Garcia-Aznar represent the ECM as a density and use experimental migration data to explore tumor spheroids [37]. Lastly, Ruscone et al. study cancer invasion using PhysiBoSS [38, 39], an implementation of PhysiCell [40] with the MaBoSS Boolean network simulator [41], producing a model incorporating intracellular signaling activated by contact with a bulk ECM [42].

Finally, there are models that explicitly represent fiber orientation or fiber orientation and density. Chauvier et al. and Hillen et al. use transport models to look at ECM [43, 44]. This work was extended by Painter to introduce a macroscopic ECM density and fiber orientation which was then applied to model tumor invasion [45]. Other authors model ECM directly as a phase in a cellular-Potts model exploring angiogenesis, tumor invasion, and impacts of cytoplasm and nucleus interaction on migration [46–48]. Some authors also form long-range fibers by viewing sets of contiguous points occupied by ECM as fibers. They can then more directly represent fiber orientation and length in addition to density [49, 50]. Park et al. use a modified Vicsek (flocking) model [51] to explore contact inhibition of locomotion in the formation of heterogeneous ECM fiber orientations [52]. Suveges et al. use a hybrid agent-based model and two phase continuous-ECM field approach (oriented, fibrous phase and non-fibrous phase) to study collective migration on oriented ECM [53]. Dallon, Sherratt and collaborators have multiple studies on ECM that include a concept of density and orientation and focus primarily on wound healing and scar formation. The models can be split into hybrid models (discrete cells and continuous ECM) or fully continuous models. Examples of the continuous models include reaction-diffusion and integro-partial differential equation models of wound healing [54, 55]. The hybrid-discrete continuous models include a model of ECM fiber reorientation, an extension to include ECM density, and a third extension to include the impacts of time varying transforming growth factor *β* profiles on all models including ECM contact guidance [56–58]. For additional details on these foundational works in modeling the ECM and wound healing, see Sherrat and Dallon’s review [59]. In a later paper, the model is further extended to combine chemotactic and ECM contact guidance, still in the context of wound healing [60]. Finally, in work similar to Dallon, Sherratt, and colleagues, Martinson et al. model a field of discrete ECM puncta (dots of immature proto-fibrous ECM) and neural crest cell-driven maturation and alignment of the puncta fibers. Using both number of ECM puncta and their orientations to alter cell motion, they study how these factors influence collective migration of neural crest cells [61].

In this work, we draw on these prior modeling approaches to capture essential aspects of bidirectional, local cell-ECM interactions. We divide the ECM into volumetric elements which track overall local ECM density, fiber orientation, and fiber-fiber alignment (anisotropy), properties collectively referred to as the local microstructure. Individual cell agents can remodel each of these properties, while these properties can influence cell behaviors including migration speed, chemotactic response, motility direction, proliferation, death, secretion, and differentiation. This mesoscale approach spans between a coarser ECM density field and more fine-grained approaches that might represent each fiber. It yields a higher dimensional characterization of ECM than ECM density alone, allowing a more diverse class of modeling assumptions, but eliminates the complexity of discrete force-based frameworks and high-order PDE models (particularly those with tensorial terms), simplifying integration with a broader class of cell behaviors than in prior works. It opens the possibility of linking important aspects of cell biology such as secretion, phagocytosis, effector attack, differentiation, proliferation, death, and cell fusion to this higher-dimensional set of ECM properties as they vary both spatially and over time. We implement this as a framework that can be readily used to incorporate local ECM effects into agent-based models, pairing it with the open source package PhysiCell [40]. To demonstrate the versatility of the framework, we present four sample models focused on cell migration and local degradation, deposition, and reorientation of matrix. In the first example, to test the framework’s ability to capture the complex effects of prior models, we replicate aspects of a study from Painter in which a front of tumor cells invade different scenarios of ECM configurations [45]. For the second example, we model wound-related fibrosis where activated fibroblasts generate a region of high ECM density that cells cannot enter or leave. In the third, recruited fibroblasts degrade a basement membrane and allow a tumor to progress from locally confined to local invasion. The final example is a model of collective migration which can produce both stigmergy (indirect coordination of migration through changes in the environment) and leader-follower migration, with more aggressive leader cells enabling mixed clusters of cells to invade surrounding tissue. Overall, our framework builds on previous work and is especially inspired by Dallon et al. [56], who developed a model of cell-ECM interactions focused on wound healing. We generalized a similar model into an extensible framework that can be rapidly adapted to new problems, in particular through use of emerging rules-based formulations [62].

We present our work as follows: Section 2 introduces the ECM representation and cell-ECM interactions and implementation details. Sections 3.1, 3.2, 3.3, 3.4 present the example models. Section 4 discusses findings, limitations, and future directions for the framework. Our supplementary material (Section 7) covers the cell-ECM interaction model in detail as well as details of cell-cell and cell-environment mechanics [40], PhysiCell rules [62], and example model parameters.

## 2 Methods: Framework description

In this section, we first provide the conceptual background and formulation of the ECM and cell-ECM interactions framework (the ECM model). Then, we illustrate the framework with a series of computational results highlighting important aspects of the framework. Finally, we give an overview of the computational details.

### 2.1 ECM and cell-ECM interaction models

To enable explorations of ECM-mediated cellular communication and interactions, we developed a three variable ECM representation (orientation, anisotropy and density) and interaction rules for cells and the ECM. We use an agent-based approach to represent cells and a continuum model to represent the fine structure of ECM. Broadly, individual ECM elements are locally remodeled, or modified, by cell migration and ECM production and degradation and cell motility, proliferation, death, and other behaviors can be impacted by ECM element variables.

#### 2.1.1 Extracellular matrix representation

Our model is intended to capture key aspects of a small region, or element, of ECM. We conceive of an element of ECM as having three variables: an average fiber orientation (*orientation*), a relative local fiber-fiber alignment correlation (*anisotropy*), and a quantity that captures the relative amount of ECM material present in an element (*density*). Assembling many of these elements together makes a whole ECM. Figure 1a shows an ECM, which is comprised of tiled ECM elements. The inset shows a single element. It is conceived of as containing many fibers (for illustrative purposes, shown as grey line segments). However, the individual fibers are unmodeled and instead an ensemble of fibers are represented through an average orientation and anisotropy (the black double-headed arrow and its length) and volumetric density (illustrated as number of fibers per element in the schematic).

**Fig. 1:**
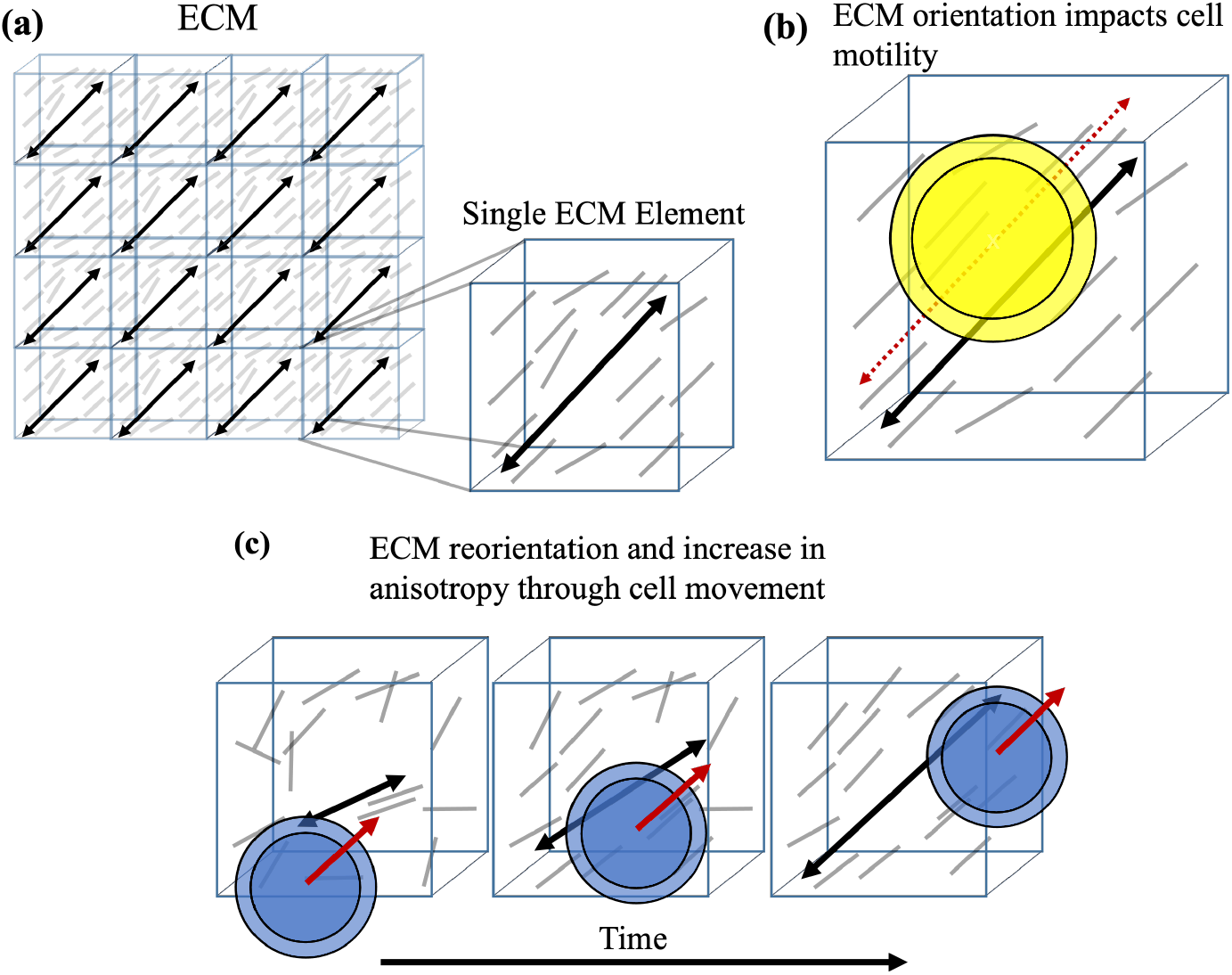
Schematic of ECM model and motility-based cell-ECM interactions. **1a** Illustration of ECM, composed of tiled ECM elements. We conceive of an element of ECM as containing many fibers. However, rather than modeling each of those fibers, we instead model the ensemble, or mean, properties of the set of fibers in a given volume. We show a schematic of a single element containing many fibers (grey line segments, for illustration purposes only) which we quantify by an overall (average) fiber orientation and anisotropy (amount of self-alignment), shown by the black double-headed arrow and its length. **1b** An element’s overall fiber orientation impacts a cell’s motility, providing a preferred angle of travel through the element (dashed red arrow). The degree of influence is related to the amount of anisotropy and the cell’s ability to sense and respond to the ECM cues. **1c** Cell movement (cell direction - red arrow) remodels ECM by aligning its element’s orientation and anisotropy. Cells align an element’s orientation parallel to their direction of travel and increase anisotropy proportional to cell speed.

Mathematically, we center an element at a location **x** in space and define the element variables as follows:

- ***Orientation:* f** (**x**), the average local fiber direction, represented as a unit length three-dimensional vector
- ***Anisotropy:*** *a*(**x**), average local fiber-fiber alignment correlation (range 0 - 1). At zero, there is no correlation and at one, locally there is complete fiber-to-fiber correlation
- ***Density:*** *ρ*(**x**), volume fraction of fibers, which we will refer to as the average local fiber density (range 0 - 1). Zero refers to complete absence of fibers and one to completely packed with fibers without void space.

Without a loss of generality, for our simulations and implementation, we arrange the elements in a Cartesian grid.

#### 2.1.2 Cell-based modeling framework

We use the PhysiCell tissue modeling system as our agent-based modeling framework [40, 62]. PhysiCell uses a center-based, off-lattice representation of cells and implements physical cell-cell mechanics. It takes a phenotypic approach to encoding cell behaviors with each cell-agent carrying its own parameters and behaviors. Additional details of the mechanics and rules used in PhysiCell are in Supplementary Material Sections 7.3 and 7.4 respectively. To model the diffusive microenvironment, we use BioFVM, which comes with PhysiCell. BioFVM is a diffusion solver that uses a Cartesian representation of space and finite volume method to efficiently handle multiple diffusing substrates over tissue-scale sized domains [63].

#### 2.1.3 Cell-ECM interactions

We also model interactions between cells and the ECM. The conceptual model is below with the detailed mathematical model provided in Supplementary Material Sections 7.1, 7.2, and 7.4.

*ECM impacts cell migration (Figure 1b):*

- Fiber orientation provides directional cues [6–12].
- Anisotropy gives a strength of directional cue: high anisotropy increases an ECM element’s influence on direction of cell migration [12].
- Density influences cell speed: too little ECM, cells have nothing to attach to; too much, cells cannot pass [4, 5].

*Cell migration and movement impact microstructure (Figure 1c):*

- Direction of cell migration reorients an ECM elements’s orientation [1–3].
- Cell-ECM element contact increases ECM anisotropy proportional to cell speed [1–3].
- Cells remodel ECM density towards a target value [1–3].

This model is motivated by findings in the developmental, disease, and tumor biology literature as well as inspired by previous modeling efforts [1, 9, 10, 56, 64]. The cell-ECM interactions are specified at the cellular level, enabling a variety of cell-ECM interactions, in particular changes in cell motility and ECM remodeling capabilities [6, 10, 64]. Additionally, ECM variables can be used to impact other cellular behaviors such as proliferation and death. Finally, we note and will demonstrate that these features can be integrated with others such as sensitivity to chemical cues and cell-cell adhesion to obtain an even richer range of cell behaviors.

### 2.2 Individual features of the cell-ECM interaction model

In this proof of concept section, results isolate and demonstrate a single or small number of aspects of the cell-ECM interaction model presented in Section 2.1. To aid in isolating only ECM-to-cell and cell-to-ECM effects, we disabled cell-cell mechanical interactions by setting cell-cell adhesion and repulsion (*C*_cca_ and *C*_ccr_ in Supplementary Material Equations 22 and 24 respectively) to zero. We used multiple cells per result to highlight the uniformity of interactions across the computational domain and as a visual aid rather than to explore how cell-cell interactions may impact the simulations.

#### 2.2.1 Cells alter microstructure, producing directional signals in the ECM

##### Combing the ECM: Local ECM fiber orientation and alignment remodeling produces extended paths in the ECM

Remodeling of the ECM elements can produce paths that ECM sensitive cells can then follow. In Figure 2a, a column of cells move deterministically from left to right across a field of un-oriented ECM aligning it in the direction of travel. Each ECM element is initialized with a random orientation and anisotropy of 0.9. Remodeling cells move across the domain, producing individual changes in ECM elements as they travel. After reaching the simulation boundary, the cells reset to the opposite boundary, and remodeling continues. In this case, after four remodeling passes (transits) through the matrix, the ECM is aligned almost completely parallel to the direction of travel with the additional effect of a slight increase in anisotropy, demonstrating ECM reorientation, and producing oriented paths across the domain. Note that we run this simulation in 2-D to aid in visualization and to demonstrate uniformity across the domain. A video is linked in the Supplementary Material (Section 7.6).

**Fig. 2:**
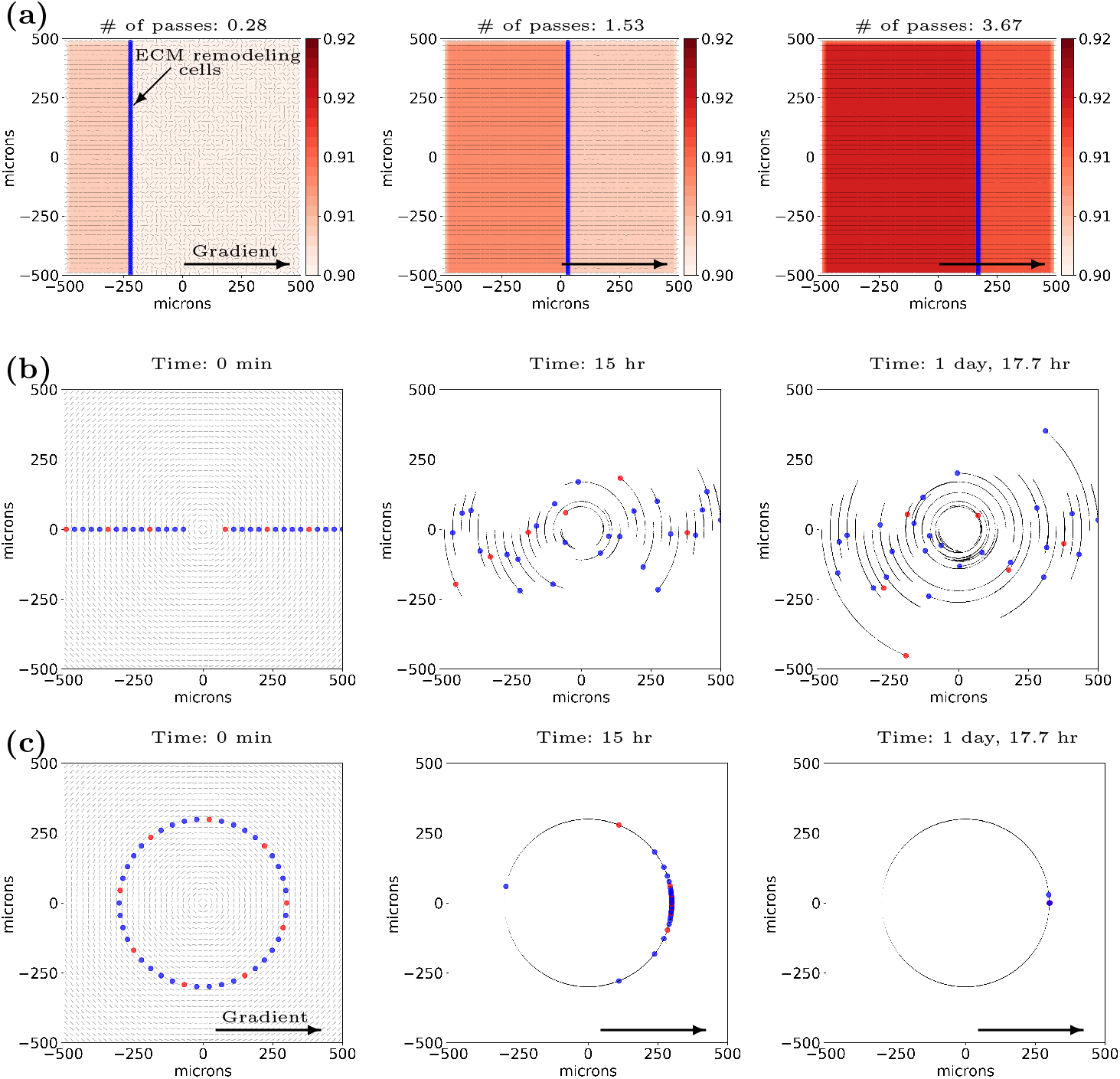
Highlights of individual aspects of the cell-ECM interactions: (**2a - Modifying ECM microstructure**) A line of ECM remodeling cells (blue circles), driven by a constant chemical gradient (black arrow), move in the positive x-direction, reorienting the initially randomly oriented ECM element orientations (black line segments) (left-most image in **2a**). Upon reaching the simulation boundary, the cells reset to the opposite boundary, and remodeling continues (middle - 1.53 passes). At four passes (left region of the right-most image in **2a**), each ECM element is almost completely reoriented parallel to the direction of travel of the cells and anisotropy has increased from 0.90 to ∼0.92. Note, we run this in 2-D to aid in visualization and demonstrate uniformity across the domain. **2b, 2c - Following ECM**. Both simulations use ECM elements that are highly anisotropic (*a* uniformly set to 1.0) and element orientations arranged to form approximate concentric circles. The simulation in **2b** begins with ECM sensitive cells almost equally spaced along the line y=0 (leftmost image - cells shown in red and blue; color is for visual contrast only). Cells follow the ECM circles in an approximate 1-D random walk due to ECM following and the lack of an external driving direction (*e*.*g*. chemotaxis), forming circular arcs (small black arrows marking cell position history every 6 minutes). The piecewise linear representation of the circular ECM tracks are more accurate as distance from the center increases (and curvature decreases). Because of this, cells following the farther piecewise linear tracks stay closer to the theoretical circular paths. The simulation in **2c** contrasts with **2b** by providing an additional environmental cue - a chemical gradient. As the ECM sensitive cells integrate both signals, in contrast to **2b**, their motion is no longer random and they travel uniformly in the direction of the gradient. With a circular initial configuration (left), the cells follow each other along the same circle (middle and right), coming to a point of relative stability where the ECM orientation is exactly perpendicular to the chemical field. Note that the ECM is static in **2b** and **2c** and does not vary across time. Videos are available. See links here.

#### 2.2.2 Cells read signals in the ECM, impacting their migration

All results in this section were produced with anisotropy set uniformly to 1.0, the maximum and most impactful value, and density uniformly set to 0.5, the value producing maximum cell speed (*ρ*_ideal_ from Supplementary Material Equation 17). Additionally, the chemical and physical (ECM) environments are static in these cases, meaning no changes in chemical substrate values or ECM remodeling occurs. This ensures that only dynamics between a cell and the ECM or respectively among a cell, ECM, and chemical field impact the results. In this set of simulations, the cells are highly sensitive to ECM (*s* = 1.0 from Supplementary Material Equation 9) as well as to the chemical gradient (*b* = 1.0 from Supplementary Material Equation 15) when present.

##### Cells can closely follow paths in the ECM

Supplementary Figure 3 shows influence of ECM on cell migration (ECM contact guidance) combined with a chemotactic field going to the right. The leftmost subfigure shows the initial cell positions as well as the ECM element fiber orientations (45^*°*^ and -45^*°*^ from the x-axis in the top and bottom sections of the domain respectively). Without ECM influence, the cells would simply move to the right. Instead, the ECM biases the cell paths away from the path set by the gradient. The middle and rightmost images include the cell’s positional history, shown as arrows trailing the blue and red cells (note in this and in all images in this section, the cell color is simply for visual contrast). A video is linked in the Supplementary Material (Section 7.6).

##### Lacking additional directional cues, cells perform approximate 1-D random motion along ECM paths

Figure 2b shows the results of cell motion on circular ECM. We show the initial cell configuration as well as the ECM element orientations on the left with the middle and right images including cell positional history at 15 hours and 41.7 hours simulated time respectively. Our ECM model does not encode a preferred direction. Instead, without an externally set direction (*e*.*g*. a chemotactic direction), cells select a random direction (see the end of Supplementary Material Section 7.1). However, contact guidance limits cell motion to be along the circular ECM paths. Thus, we observe approximate 1-D random motion along the ECM circles in this proof of concept simulation. Because of the curved tracks and error due to ECM discretization in highly curved regions (center of domain in this case), contrasting with Supplementary Figure 3, some cells move off their initial paths out to ECM circles of higher radius. We expand on this phenomenon in the discussion (Section 4). Due to the high impacts of ECM discretization at the center of the domain, for this simulation, we exclude the central region with radius of 80 *µ*m or less. A video is linked in the Supplementary Material (Section 7.6).

##### Cells can integrate environmental signals and follow non-linear paths

Figure 2c shows the results of cell motion transiting circular ECM with an additional cue supplied by a chemical gradient. This additional information causes the cells to follow the ECM circle on which they were initially placed to the right (the direction of the chemical gradient). Note that at locations where the chemical gradient and ECM orientation are perpendicular, cells are in a quasi-stable state as the two directional cues cancel. This meta-stable state is seen in the lagging cell in the middle plot of Figure 2c. Note that all other cells in this plot, except the lagging blue cell on the left side of the image, have moved many cell lengths from their initial positions. This cell began at the x-axis, which defines a line in which all units of ECM have an orientation perpendicular to the chemical field. Likewise, all the cells converge at the x-axis, where their initial ECM contour becomes perpendicular to the chemical field. Unlike in Figure 2b, with cells placed further from the origin and in a region of lower curvature, we do not observe “track jumping”. A video is linked in the Supplementary Material (Section 7.6).

### 2.3 Computational method details and data availability

This work is implemented and run in PhysiCell 1.12.0 [40]. It also uses the recently introduced PhysiCell rules [62]. All code is available with a permissive BSD license, including the ECM extensions and sample models, at GitHub with the latest release located [here](https://github.com/PhysiCell-Models/collective-invasion/releases/). The code is cross-platform capable and has been successfully compiled and executed on Ubuntu 22.04, Windows Server 2022, and MacOS 12. All simulation data for this paper is available for download at [Zenodo](https://doi.org/10.5281/zenodo.13770006). A free, interactive cloud-hosted version of the leader-follower model (see Section 3.4) is available at [nanoHUB](https://nanohub.org/tools/physicellecm) (note - sign up is required).

On consumer-grade hardware (2019 Macbook Pro), typical wall times were 13 minutes for the longer fibrosis simulation and 10 minutes for the invasive carcinoma and leader-follower simulations. The invasive cellular front had typical wall times of 3-4 minutes. On high-performance computing hardware, mean wall times were 4 minutes for the fibrosis simulation, 3.2 minutes for the basement membrane degradation and leader-follower collective migration simulations, and 1 minute for the invasive cellular front. The simulations start with 412 (fibrosis), 517 (basement membrane degradation) and 703 (leader-follower) cells and simulate 15 days, 10 days, and 10 days respectively. The fibrosis model has a decreasing number agents as the simulation progresses but also simulates 50% more time, hence its longer run time. The invasive cellular front starts with 30 cells, building to 900 cells when the simulations completes at 5 days.

We performed initial model exploration using DAPT 0.9.2. DAPT is a straight-forward way to organize model exploration on a mixed compute resources for small teams. For more information see Duggan et al. [65].

We used xml2jupyter to transform this command line C++ code into a Jupyter notebook-based graphical user interface [66]. The resulting interface is deployed at [nanoHUB](https://nanohub.org/tools/physicellecm).

## 3 Framework examples: Invasive cellular front, fibrosis, basement membrane degradation, and collective migration

We present four modeling vignettes to demonstrate framework features and ability to represent a range of biological phenomena, with a focus on ECM remodeling and cell motility. To help isolate dynamics due to physical interactions and cell motility, the presented examples do not include cell proliferation and death (except where required) as features. We use the default cell definition from PhysiCell, varying relevant parameters and behaviors as noted in each example. To examine the robustness of the simulations to stochastic variation, we ran each primary model 21 times and include a representative example for each modeling vignette.

For the chemical microenvironment, we use default settings unless noted otherwise. We pre-condition the domain by running the diffusion solver for 10 simulated minutes before starting the simulation. For ECM, we use the model described in Section 2.1. We place the ECM elements overlaying the BioFVM voxels exactly; the two meshes have a shared origin and coordinate system. See each example for exact microenvironment set up.

Finally, we note that the models were developed as 2-D simulations with cells confined to a single layer of 3-D voxels.

### 3.1 Invasive cellular front pushing into ECM

As a test of this framework’s ability to express prior complex models, we reproduced aspects of Painter’s 2009 computational study [45], in particular the first example: a model of an expanding tumor pushing cells into surrounding, structured ECM, as illustrated in Figure 10 of Painter 2009. In that work, a constant influx of cells was introduced on a domain boundary, with cells exhibiting strong ECM contact-guidance motility. The ECM orientations were either random, parallel to the domain edge, perpendicular to the domain edge, or a mix of parallel and perpendicular orientation. The work found that randomly and perpendicularly oriented matrices permitted faster invasion compared to the parallel setting and that the mixed setting produced a heterogenous pattern of invasion, with fastest invasion where the ECM was perpendicular to the interface. We tested whether the framework could be adapted to these scenarios and qualitatively reproduce their findings.

To emulate these scenarios in our agent-based system, we introduced a constant flux of cells (instantiate 30 new cells every 180 minutes along domain edge) at the bottom edge of a 600 *µ*m by 1000 *µ*m domain and simulated for 5 days (final cell count of 900). The cells had cell-cell adhesion (*C*_cca_) and repulsion strength (*C*_ccr_) of 0.4 and 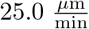 respectively, base speed *S*_max_ of 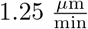, and ECM sensitivity, *s*, of 1.0. The cells were set to not remodel ECM (ECM is static) and the ECM was highly anisotropic (*a* = 1.0 throughout domain). In this scenario, cell-cell repulsion and base cell speed are increased relative to the default values of 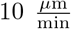 and 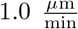. For additional parameter and simulation details, see Supplementary Material (Section 7.5.1). See Supplementary Material Section 7.3 for details on the cell-cell adhesion model. We quantified the speed of the invasive front through histograms of cell positions in the y-axis (40 bins of 25 *µ*m each) then determined the histogram bin containing the 95th percentile of cell count at five simulated days. We produced stochastic replicates for each scenario.

Consistent with the work by Painter, cells traveled further in matrices with perpendicular and random ECM orientations compared to the parallel setting. Over the 21 replicates, the fastest advances of the invasive front occurred when the ECM orientations were perpendicular to the front followed by the random orientations and finally parallel orientations. Summary statistics and visual comparison across replicates are shown in the Supplementary Materials (Supplementary Table 19 and Supplementary Figure 4). Figure 3 shows representative examples of the stochastic replicates we conducted. The changes in the invasive front across scenarios can be seen in both the deeper penetration of cells into the domain in the perpendicular and random cases as well as the cell count contours (black-white horizontal bars) in Figure 3. Furthermore, the cell number profiles (insets in upper right of the random, parallel, and perpendicular ECM scenario plots) also show differences. In the more restrictive case of parallel ECM, a relatively sharp boundary is formed, as seen in the changing slope of the cell count profile, compared to the more shallow slope of the profiles in the random and perpendicular cases. Additionally, the random orientation gives front shapes similar to a diffusive front (e.g., similar to those seen in Fisher’s equations [67]), whereas the scenario with perpendicular orientation spreads faster suggestive of superdiffusive phenomena [68, 69]. Finally, in the mixed orientation scenario, we see that invasion was fastest in the region with orientations perpendicular to the interface and slowest in the regions of parallel orientations, matching Painter 2009 [45]. See Figure 3, with links to videos available in the Supplementary Material (Section 7.6) and full stochastic replicates available in the data supplement at [Zenodo](https://doi.org/10.5281/zenodo.13770006).

**Fig. 3:**
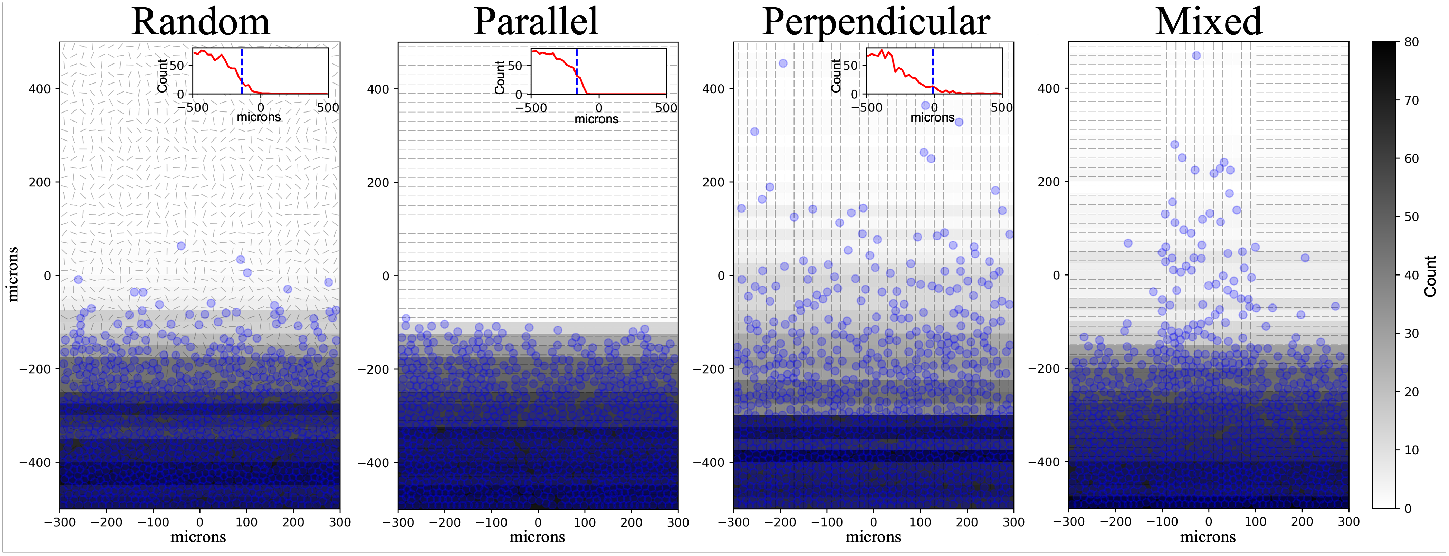
Invasive cellular front pushing into ECM: To reproduce aspects of Painter 2009’s first modeling example of an expanding tumor [45], we produced four ECM orientation configurations - random, parallel to advancing front, perpendicular to advancing front, and the mixed case of both parallel and perpendicular. In all cases, the ECM was static (no ECM remodeling) and cells entered the simulation at the bottom edge of the domain, moving with random, contact-guided motility. 30 cells were introduced every 180 minutes with no flux conditions at the boundaries. We show each simulation at five days simulated time. To assess cell penetration into the domain, we produce histograms of cell positions in the y-axis (40 bins of 25 *µ*m each). The horizontal contours are shaded according to the histogram bin counts. The insets in the upper right of the random, parallel, and perpendicular conditions show the binned cell counts viewed in profile. The dashed blue line indicates the center of the strip where 95% of total cell count is passed. ECM orientation shown in grey line segments.

### 3.2 Wound healing: ECM deposition, fibrosis, and enclosure of a wound

We produced a simple model of fibrosis. We model the recruitment of cells to clear an injury - first macrophages and subsequent recruitment of fibroblasts to rebuild the ECM in the area of the injury [17, 70, 71]. Fibroblasts are activated in the presence of macrophages, depositing ECM, leading to formation of a circle of very high ECM density and subsequent exclusion of cells from the wounded area.

#### Model results and cellular behaviors

We use three cell types - distressed/dead cells, macrophages, and fibroblasts similarly to [72]. The key rules of the model are summarized in Table 1. The distressed cells die immediately and release debris after having already begun to release debris prior to dying (Figure 4a). This is similar to an injury or some other insult that could cause a small region of cells to die at once. The debris recruit macrophages (shown in red) which chemotax up the debris gradient (not shown). Upon reaching the damaged cells, macrophages phagocytose the dead cells and also send out an inflammatory signal to initiate additional tissue repair. This signal recruits fibroblasts (yellow) via chemotaxis, which attempt to repair the damaged tissue by secreting ECM materials and increase density (yellow-to-red filled contours) above *ρ*_ideal_, the ECM density of maximum cell speed (Figure 4b,c,d). As fibroblasts and macrophages remain in contact, ECM continues to be produced. This eventually leads to the entrapment of the recruited cells. As the dead cells continue to shrink due to their death process, a cyst-like relatively cell- and ECM-free region forms, enclosed within a circle of impenetrable ECM (Figure 4e,f). See the Supplementary Material Table 7 for additional details. A video of results presented in Figure 4 is linked in the Supplementary Material (Section 7.6).

**Table 1:** Fibrosis model: principle behaviors, parameters, and systems effects. See Supplementary Material (Section 7) for parameter definitions.

| Cell type | Behavior | Key parameter(s) | Key effects |
| --- | --- | --- | --- |
| Distressed cells | Secrete debris | Debris secretion rate | Recruits macrophages to dead cells |
|  | Death | NA - cells die immediately at start of simulation | Begins fibrotic cascade |
| Macrophages | Chemotaxis up debris gradients | Bias | Enables macrophages to find dead cells |
| | ECM density response | $\rho_l, \rho_{ideal}, \rho_h$ | Cell speed is reduced as ECM density moves away from ideal value (0.5) |
|  | Consume debris | Debris uptake rate | Aids in generating gradient |
|  | Consume dead cells | Dead phagocytosis rate, phagocytosis rule saturation value and half-maximum | Enables clearance of cells |
|  | Secretion of inflammatory signal | Signal rule saturation and half-maximum values | Recruits fibroblasts to region of damaged cells |
| Fibroblasts | Chemotaxis up inflammatory gradients | Bias | Enables fibroblasts to find macrophages |
| | ECM density response | $\rho_l, \rho_{ideal}, \rho_h$ | Cell speed is reduced as ECM density moves away from ideal value (0.5) |
|  | Degrading/depositing ECM density | ECM density target value, ECM density secretion rule saturation, and half-maximum values | Can trap cells if deposited at high enough amount |

**Fig. 4:**
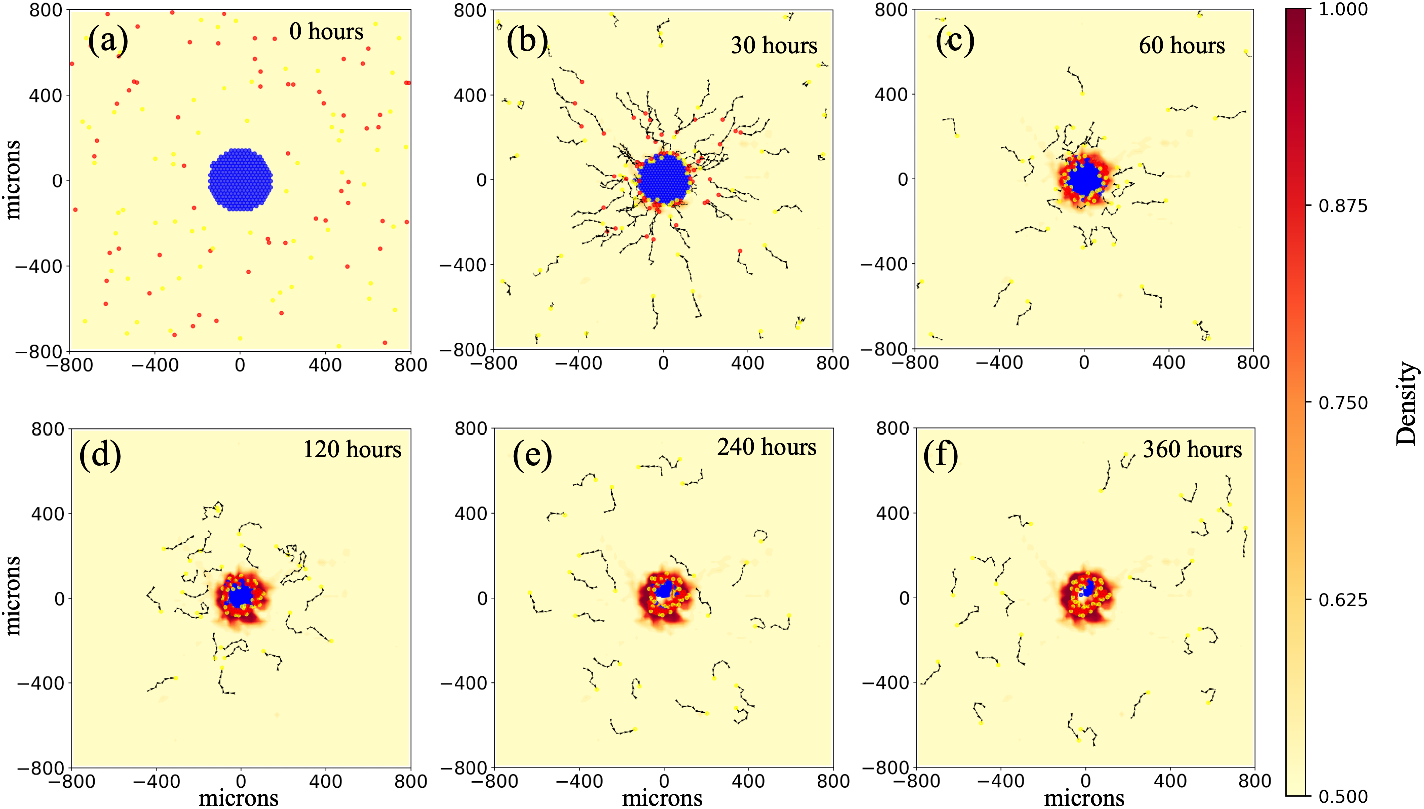
Fibroblast ECM secretion leads to cyst formation: Tissue injury, modeled as sudden death of a number of cells (blue cells) attracts macrophages (red) to remove dead cells (a, b). The macrophages recruit fibroblasts (yellow), which secrete ECM as part of the tissue repair process (c, d). However, prolonged contact with macrophages leads to continued secretion of ECM. This produces a region of impenetrable ECM which both traps the macrophages and fibroblasts and leads to a region that is relatively cell-free as the dead cells continue in their death process (e, f). Initial configuration - dead cells 175 *μ*m circle with macrophages and fibroblasts randomly placed outside of the distressed region. Yellow cells: fibroblasts. Red cells: macrophages. Blue cells: distressed/dead cells. Yellow-to-red contour plot: ECM density. Black arrows: cell positional history, going back eight time points (sampled every 30 minutes). This is available as a video. See link here.

Of the 21 stochastic replicates, 15 generated a completely enclosed region (such as the representative example in Figure 4) and six resulted in a partially enclosed region. See the simulation data set at [Zenodo](https://doi.org/10.5281/zenodo.13770006) for individual results.

#### Tissue microenvironment

We use two diffusive chemical fields in this simulation. Debris is emitted by distressed and dead cells and has a very small diffusion coefficient and no decay, representing the slow movement of cellular debris and lack of decomposition on the simulated time scales (macrophage consumption is the only removal process). This field represents the materials that distressed and dead cells release as part of the death process. We also simulate an inflammatory signal, released by macrophages as part of their response to contact with dead cells. It has a characteristic length scale of 32 *µ*m, or approximately 2.5 cell diameters. Both fields have no flux conditions.

The ECM density begins uniform with a value of 0.5 throughout the domain. Fibroblasts modify the density when recruited by macrophages. We note that in this simulation, only ECM density impacts cell motility, as the motile cell populations are modeled as being insensitive to ECM orientation cues.

See the Supplementary Material Tables 8 and 9 for additional details.

#### Tissue

Distressed cells are placed at the center of the computational domain in a circular region of radius 175 *µ*m. Cells are initialized at approximate equilibrium spacing with six neighbors (hexagonal packing). Fibroblasts and macrophages are placed randomly throughout the domain, excluding the 200 by 200 *µ*m square containing the distressed cells. Initial cell positions were held consistent across stochastic replicates. See the Supplementary Material Table 10 for additional details.

#### Comparison to previous modeling efforts

Comparing to previous work in the field, Dallon et al. 2001 studied the impacts of TGF-*β* on collagen deposition in wound healing [58]. In their model, fibroblasts deposit collagen (ECM density) in response to TGF-*β*. Comparing results between baseline TGF-*β* and increased TGF-*β*, they find more collagen deposition resulting in slower agents and less penetration of cells into the wound. While caused by a different dynamic (concentration of a signaling molecule versus contact with a different cell type), this is comparable to the slowing of the fibroblasts as they increase ECM density followed by the inability of them to penetrate fully to the center of the wound. Note also that in this simple example, we are not modeling contact guidance whereas Dallon et al. do [58].

### 3.3 Basement membrane disruption precipitating transition from local to invasive carcinoma

Here, we model the transition from a ductally-confined cancer to a locally invasive cancer, inspired by the transition of ductal carcinoma *in situ* (DCIS) to invasive carcinoma [73]. Cancer cell recruited fibroblasts degrade basement membrane (represented as high density ECM) enabling subsequent cellular invasion [9, 74, 75].

#### Model results and cellular behavior details

In this model, there are two cell types - cancer cells and fibroblasts. The key rules of the model are summarized in Table 2. In this case, we simulate the duct’s longitudinal cross section. We start with an outgrowth of cancer cells (half circle of blue agents). The cells have grown into the fluid-filled region of low ECM density - the duct lumen. They are on a band of very dense ECM (*ρ* = 1 - dark horizontal band in Figure 5a) which represents the duct’s basement membrane. The basement membrane is surrounded by stroma - with ECM density of 0.5 (Figure 5a). The cancer cells emit a tissue remodeling factor, modeled as an inflammatory factor, that attracts fibroblasts via chemotaxis with high bias (field not visualized). The fibroblasts alter the ECM and basement membrane, increasing anisotropy in the stromal region and degrading the basement membrane to a lower ECM density (*ρ*_target_ = 0.5) as they move into the tumor (Figures 5b, c, and d). Due to close proximity of fibroblasts, cancer cells alter their adhesion and affinity to each other [76], and escape, following the oxygen gradient originating at the lower simulation domain boundary (Figures 5e, f). Because the cancer cells follow ECM, they preferentially move along the groomed ECM paths, or portions of high anisotropy, but also transit the ungroomed ECM in their movement out of their location of origin (Figures 5g,i). See the Supplementary Material Table 11 for additional details. A video of results presented in Figure 5 is linked in the Supplementary Material (Section 7.6)

**Table 2:** Invasive carcinoma model: principle behaviors, parameters, and systems effects. See Supplementary Material (Section 7) for parameter definitions.

| Cell type | Behavior | Key parameter(s) | Key effects |
| --- | --- | --- | --- |
| Cancer cells | Secrete inflammatory signal | Secretion rate, target value, and secretion rule parameters | Recruits fibroblasts |
|  | ECM following | ECM sensitivity | Enables invasion of stroma |
| | ECM density response | $\rho_l, \rho_{ideal}, \rho_h$ | Cell speed is reduced as ECM density moves away from ideal value (0.5) |
|  | Chemotaxis up oxygen gradients | Bias | Enables escape |
|  | Decrease in cell-cell adhesion on contact with fibroblast | Adhesion rule saturation and half-maximum values | Enables cells to more readily escape the tumor mass |
|  | Decrease in cancer cell affinity on contact with fibroblast | Affinity rule saturation and half-maximum values | Enables cells to more readily escape the tumor mass |
| Fibroblasts | Chemotaxis up inflammatory gradients | Bias | Enables fibroblasts to find cancer cells |
| | ECM density response | $\rho_l, \rho_{ideal}, \rho_h$ | Cell speed is reduced as ECM density moves away from ideal value (0.5) |
|  | Degrading/depositing ECM density | ECM density target value | Removal of barriers to cell movement |
|  | Altering ECM orientation and fiber-fiber alignment | ECM reorientation rate and anisotropy growth rate | Provides paths for cell escape |
|  | Decrease speed on contact with cancer cells | Speed inhibition rule saturation value and half-maximum | Slows cell speed enabling remodeling to occur |

**Fig. 5:**
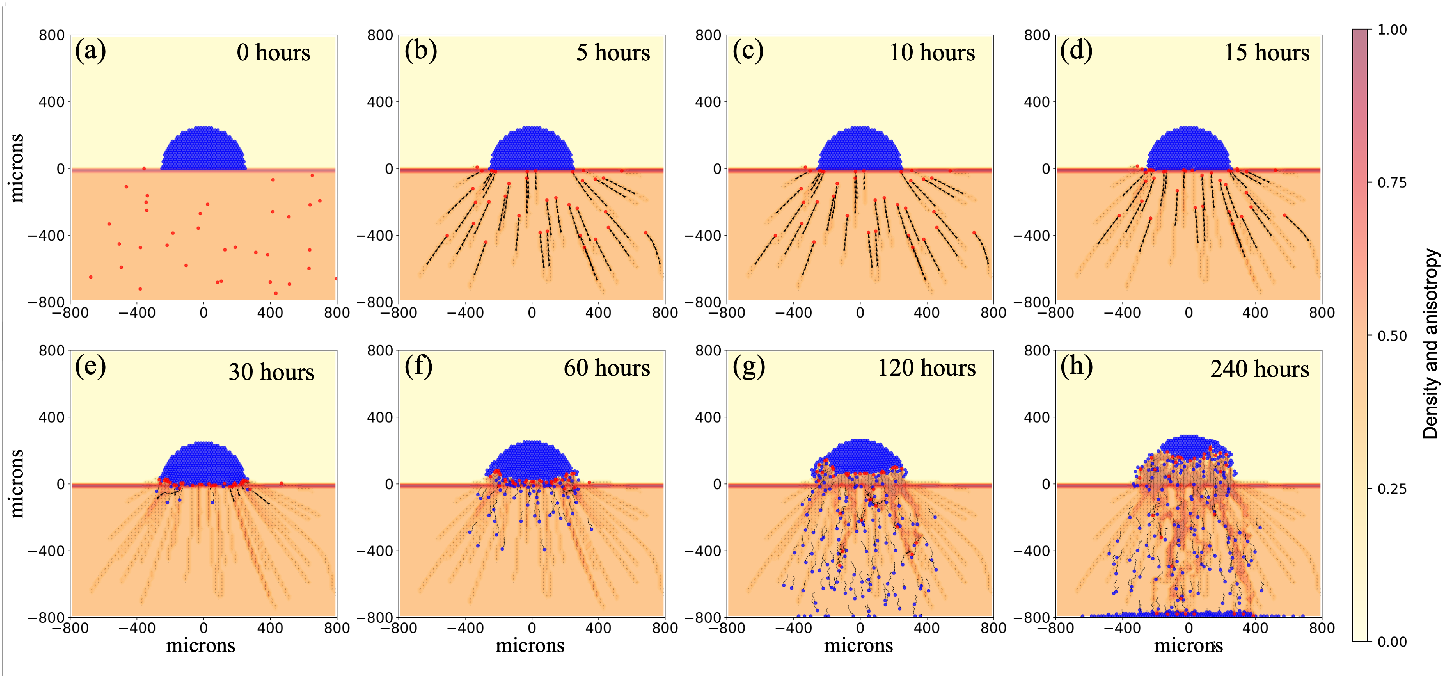
Dense ECM can form barriers while ECM degradation can lead to cell invasion: Beginning with a fluid-filled lumen, a layer of dense, impenetrable ECM, and stromal tissue (a), we see that ECM modifying fibroblasts remodel the tissue locally (b, c, d), enabling cell escape into surrounding tissue (e, f, g, i). Initial configuration - cancer cells in half circle (175 *µ*m radius) with fibroblasts randomly placed outside duct. Red cells: fibroblasts. Blue cells: cancer cells. Yellow-to-red contour plot: ECM density and anisotropy (alpha = 0.5). Black arrows: cell positional history, going back eight time points (sampled every 30 minutes). This is available as a video. See link here.

All 21 stochastic replicates generated qualitatively similar results - basement membrane degradation and subsequent invasion of stroma. Figure 5 is a representative example. See the data supplement for individual results.

#### Tissue microenvironment

We use two diffusive chemical fields in this simulation. Inflammatory signal is secreted by the cancer cells and has a diffusion length scale of 100 *µ*m in regions of high fibroblast density. We also simulate a nutrient field, modeled as oxygen (length scale of 1000 *µ*m in open tissue and 100 *µ*m in cell dense regions) coming into the duct from the bottom of the simulation. The inflammatory signal has no flux boundary conditions while the oxygen has mixed conditions: a constant value (Dirichlet condition) at the bottom of the domain and no flux conditions at the other boundaries.

The ECM begins with three zones. The top of the simulation, in which the cancer cells start, represents the interior portion of a duct (the lumen). We model it as completely fluid-filled with ECM density set to zero. The duct is surrounded by a region of very dense ECM which represents the basement membrane. Finally the duct itself is surrounded by stroma, which we model with an ECM density of 0.5. We are viewing the duct in a longitudinal cross section, rather than axially.

See the Supplementary Material Tables 12 and 13 for additional details.

#### Tissue

Cancer cells are placed in the upper half of the domain in a half circular region of radius 175 *µ*m with its main diameter in contact with the region of dense ECM (basement membrane). Cells are initialized at approximate equilibrium spacing with six neighbors (hexagonal packing). Fibroblasts are randomly placed throughout the lower portion of the domain, with placement held consistent across stochastic replicates. See the Supplementary Material Table 14 for additional details.

#### Comparison to previous literature

Comparing to previous modeling results, Kim and Othmer investigated stromal invasion enabled by basement membrane degradation [77]. In their model, also inspired by DCIS, the tumor cells themselves degrade the basement membrane by secreting proteolytic enzymes. In their model, as in ours, the breakdown of the ECM surrounding the mass of cells led to cellular invasion. In their case, they observed that a front of leading cells form as the basement membrane was broken down compared to our more dispersed breakdown of basement membrane and invasion. However, they had a point source of attractant compared to our line source. Furthermore, our basement membrane is broken in several locations and our tumor cells follow contact guidance. Both of these features tend to promote a spreading of the tumor cells compared to Kim and Othmer’s scenarios.

### 3.4 ECM contact guidance enables leader-follower migration

To address questions centered on the mechanisms of multicellular invasion, we developed a tissue simulation focusing on collective cell migration. It is inspired by the cancer biology literature [64, 78] and builds off previous cell-based spheroid modeling work [40]. Mimicking Cheung et al. [64], we include two cell phenotypes: more aggressive leader cells and less aggressive follower cells. Their observations suggest some form of communication between the leaders and followers; we model that communication as signals written in and read from ECM microstructure. We use our cell-ECM interaction model with leaders writing signals (remodeling microstructure) and followers reading those signals (contact guidance). Model details follow as well as more detailed accounting of the simulations that lead up to collective migration.

#### 3.4.1 Leader-follower model details

##### Cellular modeling

We model two different phenotypes - leader cells and follower cells. In our example, these cells represent invasive and non-invasive cancer cells, respectively, but they could represent other types of cells that exhibit similar behaviors (more motile and less motile sub-phenotypes). The key rules of the leader-follower model are summarized in Table 3. In particular, leader cells follow oxygen gradients with a high chemotactic bias and encode their movements via changes to ECM microstructure, specifically altering ECM element orientation and anisotropy (see Section 7.1 for mathematical details). Followers’ movements are influenced by signals in the ECM (contact guidance) and are chemotactic with bias equal to local value of *a* (Chemotaxis Model II - Equation 16) following previous works that used chemotactic bias [60, 61]. They read signals in the ECM with a high sensitivity (*s* = 1.0) and are incapable of remodeling ECM. Thus, in highly anisotropic ECM, followers use contact guidance and chemotax, while in unaligned ECM, they move randomly. Combining leader and follower behavior together, leaders effectively signal paths which followers read through contact guidance. See the Supplementary Material Table 15 for additional parameter values.

**Table 3:** Cell-based model: principle behaviors, parameters, and systems effects. See Supplementary Material (Section 7) for parameter definitions.

| Phenotype | Behavior | Key parameter(s) | Key effects |
| --- | --- | --- | --- |
| Leader cells | Altering ECM orientation and fiber-fiber alignment | ECM reorientation rate and anisotropy growth rate | Produces signals in ECM, providing paths for cell escape |
|  | Chemotaxis | Bias | Enables production of coherent paths |
|  | Cell-cell adhesion and repulsion | Ratio of adhesion strength to repulsion strength | Enables collective cell migration |
| Follower cells | ECM following | ECM sensitivity | Allows cells to read signals in the ECM |
|  | Chemotaxis | Local anisotropy | Enables coherent migration |
|  | Cell-cell adhesion and repulsion | Ratio of adhesion strength to repulsion strength | Enables collective cell migration |
|  | Random motion | Local anisotropy | Enables finding of paths in the ECM |

##### Tissue microenvironment

We simulate avascular tissue with a diffusing oxygen-like field coming from the domain boundaries (Dirichlet conditions). Cells uptake oxygen and there is decay in the cellular milieu. As in the basement membrane example, we use a diffusion length scale of 100 *µ*m for regions of high cell density. See Supplementary Material Table 16 for additional details.

The ECM is initialized uniformly with anisotropy of 0.0, ECM density of 0.5 (no impact on cell speed), and random fiber orientations. See Supplementary Material Table 17 for additional details.

##### Tissue

Cells are placed at the center of the computational domain in a circular region of radius 175 *µ*m and were randomly assigned to be either followers or leaders (95% followers, 5% leaders). Cells are initialized at equilibrium spacing with six neighbors (hexagonal packing), with placement held constant across stochastic replicates. See the Supplementary Material Table 18 for additional details.

#### 3.4.2 Leader-follower results

##### Signal generation and reading are required for collective behavior

We begin the exploration as a proof of concept with simplified dynamics: leader cells exhibit the limit or extreme behavior of instant ECM microstructure remodeling, using Supplementary Material Equations 1 and 2. (The previous examples, Sections 3.2 and 3.3, used continuous or non-instant remodeling - Supplementary Material Equations 3 and 5.) We use these simplified dynamics to aid exploration, not to suggest that cells necessarily modify ECM much faster than they move. Additionally, to show there is impact from the cell-ECM interactions, we isolate two model behaviors, the ability to write (matrix remodeling) and read signals (contact guidance), while disabling cell-cell adhesion. In Figure 6, we observe similar results with ECM writing only (left column: follower ECM sensitivity *s* set to zero) and ECM reading only (middle column: leader remodeling rates set to zero). In both cases, leaders separate from the initial disc of cells and reach the simulation boundary at approximately 16 hours while followers remain roughly in the center of the domain. Only with both writing and reading (right column) were most cells able to escape the domain center and invade their surroundings. The guidance provided by the ECM combined with the chemotactic behavior of followers on groomed ECM enables them to follow the leaders’ paths out of the center of the domain. This results in stigmergy, wherein agents alter their environment and in doing so, effectively signal preferred paths to other agents. Links to simulation videos are in Section 7.6.

**Fig. 6:**
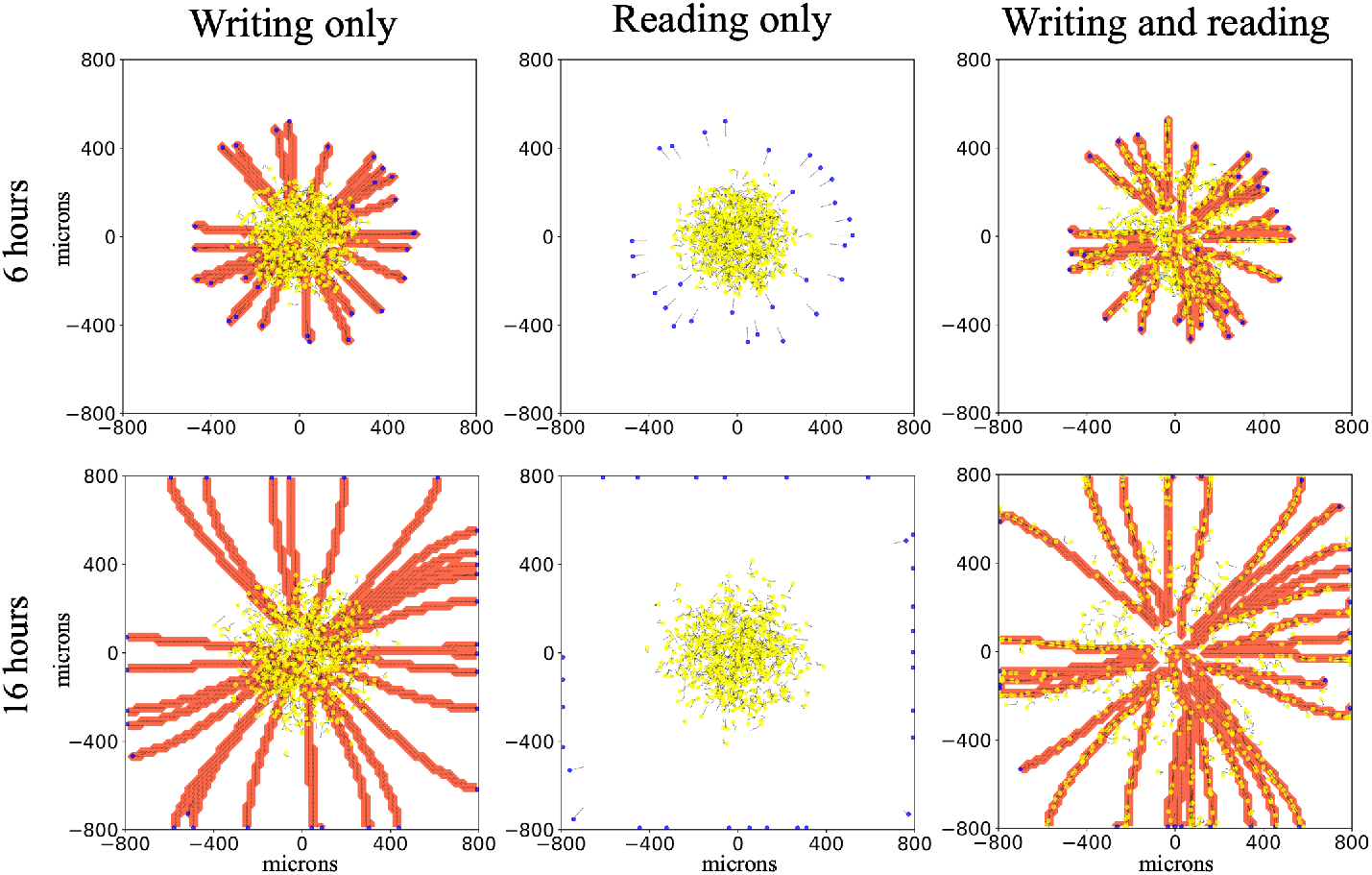
Communication of cell path through environmental modification produces stigmergic cell behavior: Modification of the environment (left) and reading of the environmental signals (middle) separately do not produce collective behavior. With both together (right column) we observe that most cells escape the domain center and invade their surroundings through the combined effects of following leaders’ paths (stigmergy) and chemotaxis. Writing only: *s*, ECM sensitivity, for followers set to zero. Reading only: 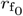 and 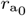, reorientation and realignment rates, for leaders set to zero. Initial configuration - 703 agents (5 % leaders) in 175 *µ*m circle. Yellow agents: followers. Blue agents: leaders. Red contour plot: ECM anisotropy - uniformly equal to one in remodeled ECM elements. Line segments: ECM orientation. Black arrows: cell positional history, going back 12 time points (sampled every 6 minutes). Videos are available. See links here.

Note that we hold the initial configuration of each simulation variation constant to isolate motility effects only.

##### Cell-cell adhesion enables collective migration

Adding cell-cell adhesion (see Supplementary Material Section 7.3 for details on the cell-cell adhesion model) to provide more realism to the simulation, while still using the limit assumption of instant signal writing, we observed a range of dynamics and cellular patterning, including collective migration in which phenotypically heterogeneous cells gather in clusters and move together while roughly maintaining the same composition over time. We see that ECM-mediated communication can still produce stigmergy (Figure 7a) when speed is high enough to overcome cell-cell adhesion. In Figure 7b, we see leaders migrated to the leading edge of cell clusters. Due to a balance between cell motility and cell-cell adhesion, cells stay in contact with each other, producing leader-follower collective invasion. Finally, when cell speed is even lower, adhesion takes over cell mechanics and relative cell positions are constant over the computational experiment (Figure 7c) while there is a small overall displacement of the whole cell mass (see video). Links to simulation videos are in Section 7.6. Note that cell-cell adhesion is of equal strength across all possible combinations of interactions (follower-follower, follower-leader, and leader-leader). Also, due to our interests in highlighting cell-ECM interactions, we have excluded exploring other possible ways to enable collective migration such as cell-cell repulsion with cell proliferation.

**Fig. 7:**
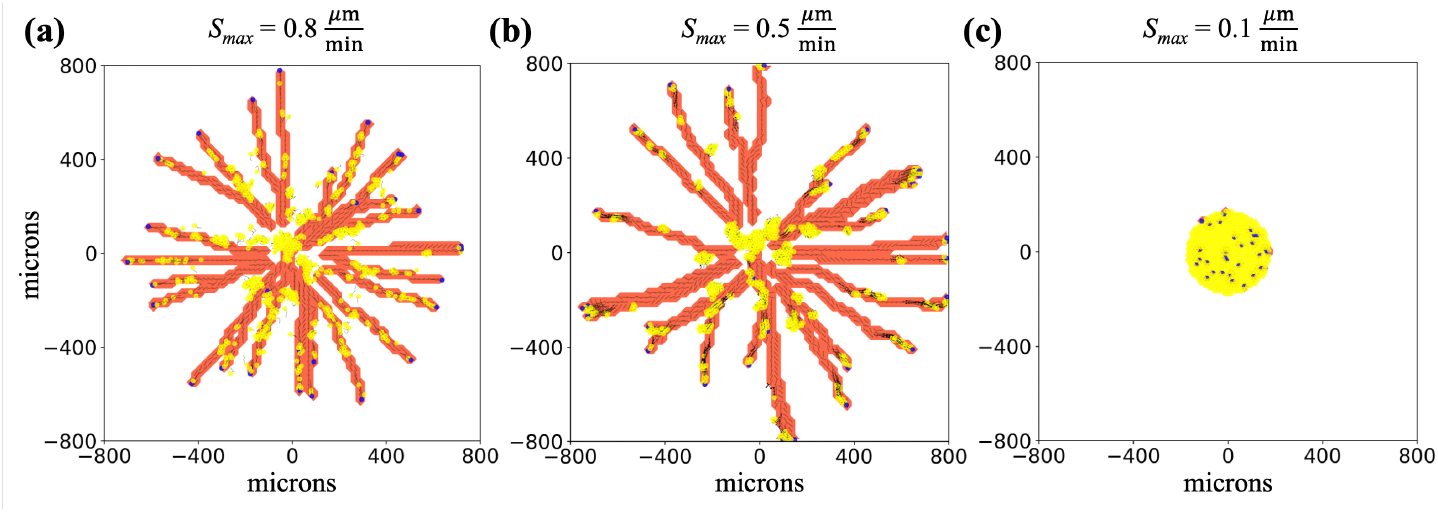
Multiple behaviors are produced with the leader-follower model by including cell-cell adhesion: With cell-cell adhesion on 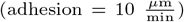 across all cell-cell interactions), multiple behavioral regimes are observed in the limit case of instant remodeling. In **(a)** with *S*_max_ (maximum cell migration speed) at 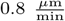, we see stigmergy with followers (yellow cells) following trails left by leaders (blue cells), as in Figure 6. **(b)** Decreasing *S*_max_ to 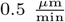 yields collective migration. In **(c)** with *S*_max_ at 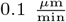, cell-cell adhesion cannot be overcome and the cell positions are frozen in the initial configuration. Initial configuration - 703 agents (5 % leaders) in 175 *µ*m circle. Adhesion set to 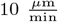. Yellow agents: followers. Blue agents: leaders. Red contour plot: ECM anisotropy - uniformly equal to one in remodeled ECM elements. Line segments: ECM orientation. Black arrows: cell positional history, going back 12 time points (sampled every 6 minutes in **a**, every 30 minutes in **b**, and every 120 minutes in **c**). Videos are available. See links here.

##### Collective migration is retained in the context of non-instant signal generation

Having shown that the cell-ECM model can produce a range of behaviors in the limit of instant signal generation, we now relax this assumption and let signal generation occur over a period of time instead of instantly. We give ECM reorientation and anisotropy non-instant rates of change and update them with Supplementary Material Equations 3 and 5, as in the previous example models. See Section 7.1 for additional details. Figure 8 contrasts the results of the limit of instant signal writing with non-instant signal writing. Figure 8a shows two time points using the limit model, with cell speed of 0.5 *µ*m/min and adhesion value of 10 *µ*m/min (the same parameters as Figure 7b). In the earlier time point (top), we see what was the initial cluster of cells broken into several smaller clusters moving out from the center of the domain. In the next time point, we see that heterogenous clusters have continued towards the simulation boundary (with some reaching it) while clusters of followers remain in the middle. In Figure 8b, relaxing instantaneous signal writing, we observe that collective migration still occurs. Leaders alter ECM with the amount of microstructure change, and thus signal strength, depending on cellular residence time, speed, and density, leading to a range of anisotropy values. Leaders remain in contact with followers and lead them out of the center of the domain. However, because altering ECM microstructure (in particular increasing anisotropy) is no longer instant, the signals to followers are not as strong compared to 8a. We observe that fewer cells are at the periphery of the domain when remodeling is slower (finite). The relative lack of strong signals in the non-instant scenario leads to less chemotactically biased migration (more random migration) in the follower population, decreasing the uniformity of velocities across a cluster of cells. Thus, as the cells adhere to each other but attempt to move in divergent directions, overall cell speed is effectively reduced (see Supplementary Material Equation 21). The more uniform the cell velocities, the closer to *S*_max_ the average speed of a cluster will be. Overall, the lack of uniformity leads to less coherent migration in which the consistently chemotactically-driven leaders break away and are then able to proceed ahead of the followers. For more details see Section 7.3 and Equation 19. The remaining followers then continue out behind the leaders via stigmergic behavior. See Supplementary Section 7.6 for links to videos.

**Fig. 8:**
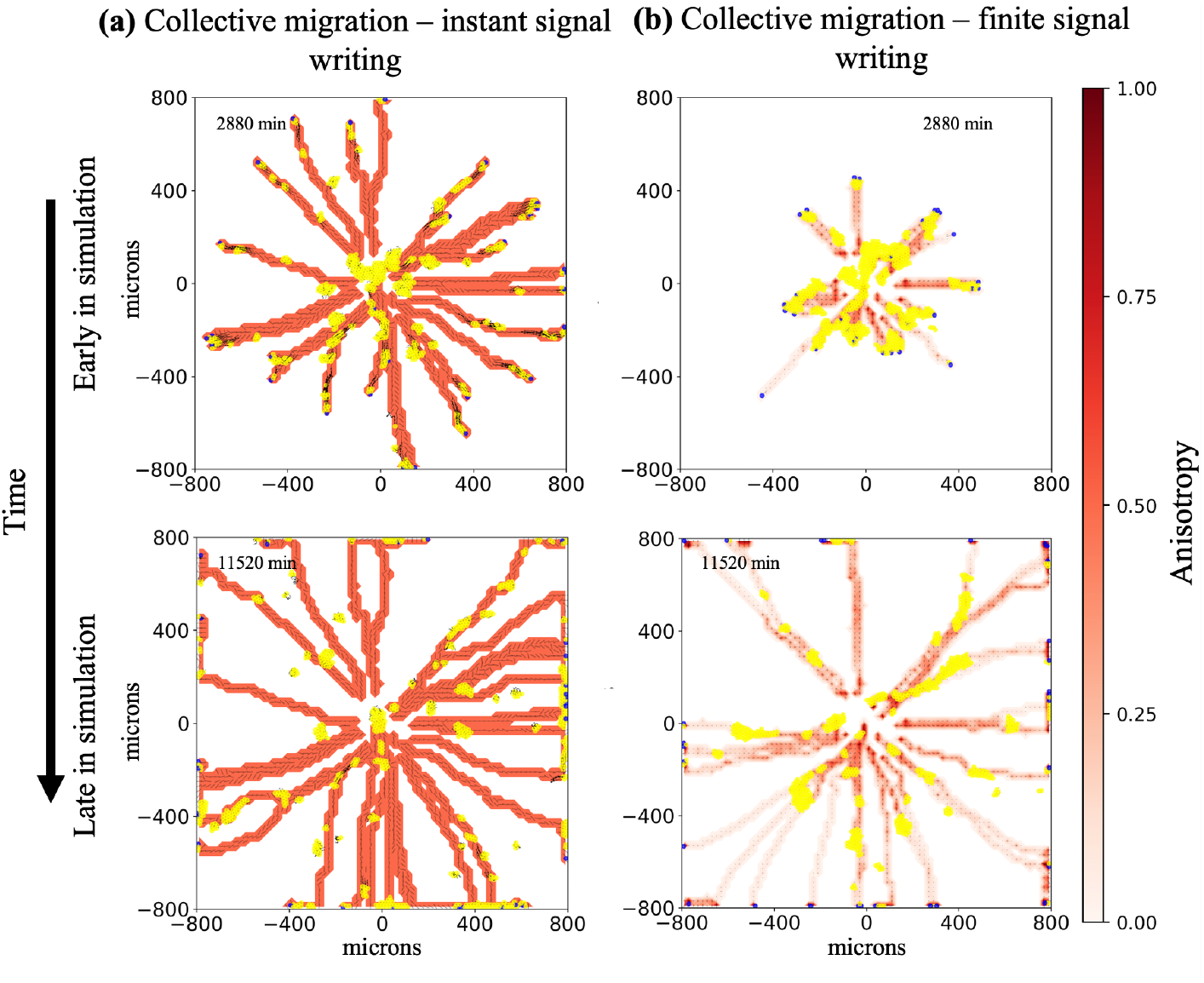
Collective migration is retained in the context of non-instant signal writing: In **(a)**, which is the limit case of instant signal generation (same parameter settings as Fig 7**b**), we observe mixed clumps of both leader (blue) and follower (yellow) cells migrating together (leader-follower collective migration). In 8**b**, we relax instant remodeling and still observe leader-follower collective migration: leaders travel at the front of mixed clusters of leader and follower cells, leading them out of the center of the domain. They do not stay in contact as long as in 8**a** but strong enough paths are generated to enable followers to continue out via stigmergy. Fiber realignment rate: 0.004 min^−1^, fiber reorientation rate: 4 min^−1^. Initial configuration - 703 agents (5 % leaders) in 175 *µ*m circle. Adhesion set to 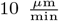. Yellow agents: followers. Blue agents: leaders. Contour plots: ECM anisotropy - (8**a**) uniformly equal to one in remodeled ECM elements and (8**b**) color bar. Line segments: ECM orientation. Black arrows: cell positional history, going back 12 time points (sampled every 6 minutes). Videos are available. See links here.

We performed stochastic replicates on the continuous remodeling collective migration simulation of which Figure 8b is a representative example. All 21 stochastic replicates generated similar results - leader-follower type invasion followed by continued stigmergic movement of followers along the leader generated ECM paths. See the data repository at [Zenodo](https://doi.org/10.5281/zenodo.13770006) for individual results.

We observed that collective migration behavior in the finite signal writing scenario is sensitive to remodeling parameters and is lost with a less than order of magnitude change in fiber realignment and reorientation rates as seen in Supplementary Material Figure 5. Fiber reorientation and realignment rates varying by a factor of four are enough to change the behavior. This model could be further tuned, matching to data related to cell speeds and patterning, to suggest conditions under which collective leader-follower migration may occur in non-computational model systems.

Examining Figures 6 and 7, in Figure 6 (without cell-cell adhesion), follower cells engage in stigmergy (right panel) while frequently straying from leader-cell created paths. In contrast, in Figure 7a (with cell-cell adhesion), straying is less frequent, implying that cell-cell adhesion enhances stigmergy, an observation deserving further study.

This model could also explore “wisdom-of-the-crowd” effects in noisily oriented ECM. Prior work has shown that the cell clusters average out noise in chemotactic gradients better than single cells [79]. Additionally, cell-ECM sensitivity, ECM anisotropy, and cell-cell adhesion could be varied to study impacts of other aspects of cell-ECM and cell-cell interactions on these effects.

Finally, we note that in this example model, we did not explore asymmetric rates of cell-cell attachment. Spring-like attachments with different rates of attachment versus detachment may better mimic cell-cell adhesion [80]. This enhanced adhesion might enable leaders to better pull cells and produce collective migration, perhaps eliminating the need for the followers’ oxygen-driven directional cue.

##### Comparison to other works

Comparing to previous biological and modeling work in the field, Ilina et al. studied collective migration and cancer metastasis, attempting to understand connections between cell-cell adhesion, cell density, and ECM confinement[81]. Their *in vitro* and *in vivo* experimental results showed that collective migration occurs with high cell-cell adhesion but also with low cell-cell adhesion (for example with knocked down E-cadherin expression) in the context of high ECM density, which corresponds to relatively decreased speed. Furthermore, the phenomena of collective behavior was investigated with an agent-based model of cells transiting along a collagen ECM and plastic culture dish interface. In particular, the authors found that low cell-cell adhesion and low ECM density, which led to higher cell speed, were associated with individual cell migration, while high ECM density (lower cell speed due to ECM confinement) or sufficiently high cell-cell adhesion both enabled collective migration. The high ECM density/low cell-cell adhesion lacked local cell to cell velocity correlations compared to the low ECM density/high cell-cell adhesion. They suggest that the system’s jamming-unjamming phase transitions, which describe the shift between a solid-like state (cells are jammed and largely immobile) and a fluid-like state (cells are unjammed and can move individually or collectively), may have several phases. These include a jammed phase, two mobile phases corresponding to the two collective migration regimes (one with low cell-cell adhesion and another with high cell-cell adhesion), and finally an unjammed gas-like phase. While we did not look at ECM-based cell confinement, this bears similarity to our results in Figure 7, with the gas phase corresponding to 7a (ECM of sufficiently low density to not confine cells), 7b corresponding to the active nematic regime, and 7c being jammed, but in this case strictly due to cell-cell adhesion. Following up on Ilina et al., Kang et al. studied collective cell migration in tumor spheroids with a hybrid vertex-particle-based model, culminating in a phase diagram of cell jamming [82]. They found that in the regime of low cell speed and higher ECM density (more regions of excluded volume within the domain), cells remained clumped together (solid phase) until a sufficient speed was reached to enable collective movement (liquid phase). Finally, with lower ECM density (relatively less volume exclusion for cells) and higher cell speeds a gas-like phase was observed. Again, these results are similar to our results in Figure 7 with the gas phase corresponding to Figure 7a, liquid phase to 7b and solid to 7c. Interestingly, both [81] and [82] either hint at or produce phase diagrams. Future work with this example model could explore the transitions between the subplots of Figure 7 as a phase diagram, in addition to considering aspects of cell confinement through use of a non-uniform initialization of the ECM and the possible role of heterogenous local chemotactic gradients in the formation of ECM paths.

Finally, while the collective migration example was inspired by the cancer biology literature [64, 78], Martinson et al. [61] produced a similar model. In this work, they model neural crest cell invasion, also using ECM remodeling cells to lead clusters of ECM following cells into tissue. While varying in some aspects (such as using haptotaxis and only cell-cell repulsion), their use of a similar leader-follower model in developmental biology hints at this model’s and the framework’s applicability to more general leader-follower migration models.

## 4 Discussion

In this work, we presented a framework designed to capture salient aspects of bidirectional local cell-ECM interactions using a relatively simple mathematical model. Drawing on previous works, we developed a model of extracellular matrix using a mesh of voxels (ECM elements), each described by a limited set of features: overall anisotropy, overall orientation, and overall density. We added cell-ECM interactions including ECM-mediated changes in cellular motility (tunable contact guidance and ECM dependent cell migration speed) and cell-mediated local microstructure remodeling (changes to ECM element anisotropy, overall fiber orientation and density). Furthermore, through the use of PhysiCell rules [62], additional bidirectional interactions are possible without the need to write, compile, and test custom interaction code.

We demonstrated the framework with four examples that focus on cell motility-ECM interactions: an invasive cellular front, wound healing, basement membrane degradation, and collective migration. Each of these examples uses the same core framework with changes in source code only for simulation initialization. We make this newly implemented mesoscale, bidirectional cell-ECM interaction framework freely accessible to the community aiming to encourage and enable sharing of expertise among multiple problem domains (e.g., tumor biology, developmental biology, wound healing, angiogenesis, fibrosis). Furthermore, our approach opens up rich explorations of cell-matrix biology by allowing direct links between a broad array of cell behaviors and ECM variables, without requiring additional coding. This represents a key methodological advance in this work: enabling the exploration of a large set of relationships between cell behaviors and ECM state.

With this base framework constructed, there are opportunities for extensions. Future models could initialize the ECM variables from high-resolution ECM data based upon analysis of the mean and variation in fiber angles (for orientation and anisotropy) and amount of material present (for fiber density). We note that our framework currently captures local interactions between single ECM elements and single cells; it does not incorporate larger-scale interactions. This contrasts with other frameworks and models that represent the network like connections within a spatially-coupled ECM or distribute cellular remodeling and contact guidance among multiple ECM elements (e.g. - [25, 29, 56]). To address non-local remodeling of ECM, future efforts will focus on incorporating the option for a continuous, force-based ECM model, for example as a visco-poroelastic material using a finite element approach, then sampling the variables established in this work on the ECM element grid. To address the current limit of remodeling only a single ECM element, we aim to explore approaches to spread remodeling over multiple, adjacent ECM elements, for example with Gaussian smoothing or other techniques as well as enabling ECM influence on cell behavior from multiple ECM elements [56, 61]. Similarly, we aim to extend bidirectional cell-ECM interactions to allow cells spanning multiple elements to influence and be influenced by each element.

As noted in Section 2.2.2, due to ECM discretization, cells may miss ECM cues in regions of rapidly changing ECM orientation (high amounts of path curvature) as observed in Figure 2b. ECM following is a function of cell speed, curvature in an ECM path, and element size. To reduce the effect of the discretization, as needed, ECM element size can be decreased by modelers noting that decreasing element size will be balanced by decreasing the area of influence in the bidirectional cell-ECM interaction and will increase computational costs. The current default size of 20 *µ*m per side was selected to be on the order of the size of the cell, as in a previous work [40]. This enables cells to modify a volume about equal to their volume, forming paths about one cell width in diameter (with our cell diameter of ≈ 17 *µ*m). Future versions of the framework that include spreading the bidirectional cell-ECM influence over a region (discussed above) might ameliorate discretization effects by effectively smoothing the path.

In the initial version of the framework, we aimed to have simple constitutive relationships that produced a range of emergent behaviors showing the utility of the underlying framework variables and relationships. As an example, we model the anisotropy as only stable or increasing. This limits the ability to capture multiple, extensive remodelings of an ECM element. Future extensions to the framework could model both growth and decline in fiber-fiber alignment. Another possible extension is stochastic contact guidance where cell direction is sampled from a distribution rather than set deterministically. This approach was recently implemented using the von Mises distribution [61], showing its feasibility and interest within the community. Moreover, in future releases, we plan to generalize the framework architecture to enable modelers to easily replace built in constitutive relationships with their own relationships. Furthermore, architecture that allows easy exchange of one constitutive relationship for another would enable useful comparisons among model relationships. Future work could include comparing simulation outcomes, numerics, and computational performance of cell-ECM interaction formulations as similarly investigated for cell-cell mechanics constitutive relationships [83].

Finally, we note that the sample models were developed as 2-D simulations (single slice of a 3-D environment), and could be, by reasonable extension of the cell-ECM interaction models to 3-D, be extended to 3-D. Also, while we have demonstrated this framework for a generalized ECM or single ECM component, the framework could readily be expanded to support a vector of ECM components. These extensions are additional future work for the framework.

## 6 Acknowledgements

We would like to thank Margherita Botticelli, Michael Getz, Adrianne Jenner, Heber Rocha, and Jordan Rozum for their expertise and assistance with this work. We gratefully acknowledge access to FutureSystems at the Digital Science Center, Luddy School of Informatics, Computing, and Engineering. This research was supported in part by Lilly Endowment, Inc., through its support for the Indiana University Pervasive Technology Institute.

## 7 Supplementary Material

### 7.1 Cell remodeling of ECM microstructure

Here we detail the mathematical models for updating ECM element variables anisotropy, orientation, and density. The ECM elements and cells are conceived of as being in close contact; cells alter the ECM element they are physically in. To formalize which ECM element a cell modifies out of the many possible options (see Figure 1a), we define cells as interacting with whichever element their center coordinates are the closest to, as in [40]. To do this, suppose cell *i* is at position **x**_*i*_. We remodel the ECM element whose center 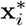 is closest to **x**_*i*_.

We start with outlining two related methods for altering ECM microstructure - the limit case of instantaneous remodeling and the non-instantaneous case.

#### Instant anisotropy and orientation remodeling

To explore an extreme case of ECM microstructure modification/ECM mediated communication, we model anisotropy and ECM element orientation as being immediately updated when contacted by a remodeling cell, resulting in the following relationships to update *a* and **f** :

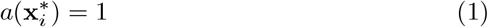

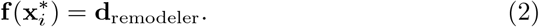

Because of the underlying updates to **f** (see next subsection), this permanently writes with high signal strength the motility direction of the first cell to pass through a particular ECM element.

#### Non-instantaneous microstructure remodeling

##### Anisotropy

Anisotropy is represented by a scalar ranging from 0 to 1, representing “complete lack of anisotropy” (or a disordered state) to “complete anisotropy” (a highly ordered, oriented state). The rate of change in anisotropy over time is given as:

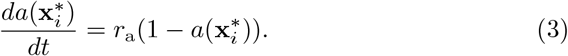

*r*_a_ is specified as:

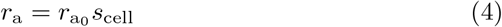

where 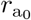 is a base rate of change and *s*_cell_ is the migration speed of the cell changing the ECM element’s anisotropy.

##### Orientation

We model changes to fiber orientation as follows:

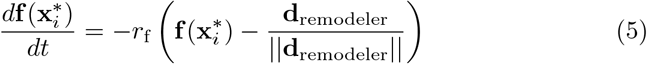

with

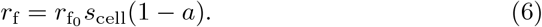

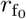 is the base rate of change, **d**_remodeler_ is the direction of the remodeling cell, and *s*_cell_ is cell speed. **f** is normalized after each update. Note that when *a* = 1, orientation cannot change. Also, we select to remodel **f** such that it takes on the smallest angle between it and **d**_remodeler_.

#### Density

Density ranges from 0 to 1, representing a region lacking ECM material to a very dense region completely filled with ECM material, respectively. Cells change ECM density locally through the following relationship:

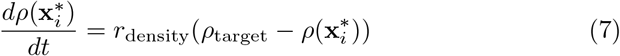

where *r*_density_ is the cell’s characteristic rate of deposition/degradation of ECM and *ρ*_target_ is the cell’s target for ECM density. This parallels chemical substrate secretion by cells in PhysiCell [40]

### 7.2 ECM microstructure influence on cell motility

Cell motion, in particular, cell speed and direction, can be altered via cell-ECM interactions. ECM element orientation and anisotropy are combined with a cell’s preferred direction to produce the cell motility vector. Independently, cell speed is influenced by ECM density enabling ECM to block or guide cell paths. We lay out the orientation and anisotropy influences first, then the influence of density. As with remodeling ECM microstructure, we define cells to be affected by and read whichever element their center coordinates are nearest as in [40].

#### 7.2.1 ECM orientation and anisotropy

We developed a method to calculate **d**_actual_, a cell’s actual motility direction, based on the cell’s preferred direction **d**_preferred_, a cell specific ECM sensitivity *s*, and local anisotropy and orientation. As illustrated in SM Figure 1, we want to constrain cell motion to the sector spanning the ECM orientation (**f**) and the cell’s intended direction of travel (**d**_preferred_) in a way that does not bias direction of travel along the fibers. We use **f** and **d**_preferred_ to form an orthonormal basis set that spans the cell’s potential directions (the sector produced by the angle *θ* in Figure 1d). We calculate **d**_actual_ by blending the basis vectors using the cell’s ECM sensitivity parameter *s* and local ECM element’s anisotropy *a* as follows:

**Supplementary Figure 1:**
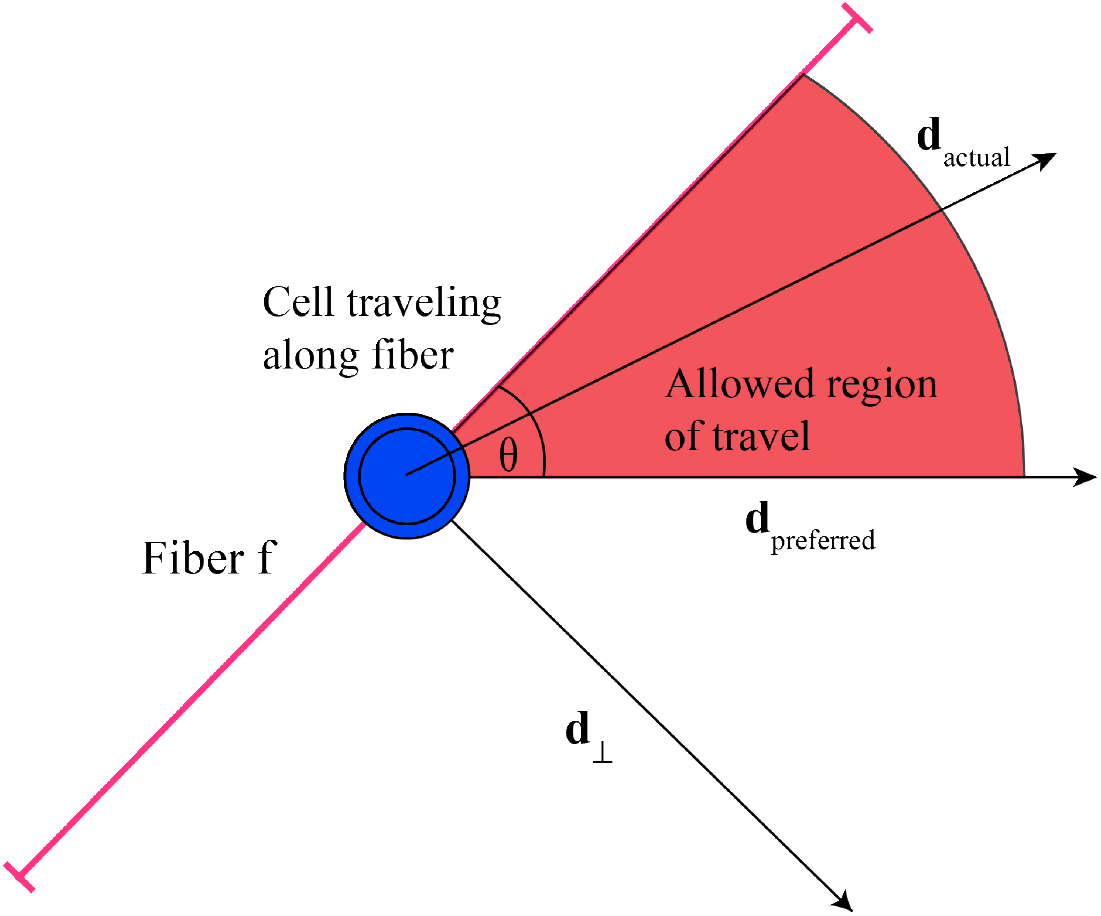
Schematic for mathematical model of cell motility and ECM orientation interactions. Cell shown in blue. See 8, 10, and 12 for variable explanations.

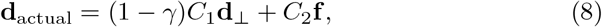

where *γ* is the effective ECM influence parameter. This captures both cell sensitivity and magnitude of ECM fiber alignment (signal strength):

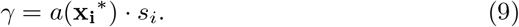

*s*_*i*_ and 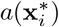 are the sensitivity of cell *i* and anisotropy at that location respectively. The other terms in SM Equation 8, the coefficients *C*_1_ and *C*_2_ and basis vector **d**_⊥_, are calculated by decomposing **d**_preferred_ and basic trigonometry. For brevity, we will drop location and indexing of cells unless needed for clarity.

We find **d**_⊥_, using the trigonometric relationship **A** *·* **B** = ||**A**||||**B**||cos *θ*. Letting **f** and **d**_preferred_ be unit vectors and *θ* being the angle between them, the relationship simplifies to:

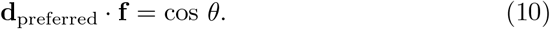

Scaling **f** by that dot product, we get the component of **d**_preferred_ that points along **f**. Subtracting that result from **d**_preferred_, we obtain a vector perpendicular to **f** as follows:

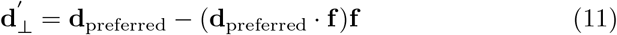

with normalization producing the unit vector **d**_⊥_ seen in SM Equation 8 and SM Figure 1.

Note also that **d**_preferred_ can be decomposed as follows:

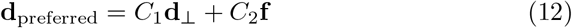

with *C*_1_ and *C*_2_ the same as in SM Equation 8. They are determined via dot product and again, we take advantage of the terms being unit vectors:

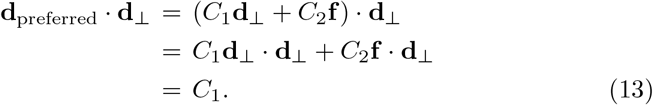

Likewise, dotting both sides of SM Equation 12 by **f**, we obtain *C*_2_ as follows:

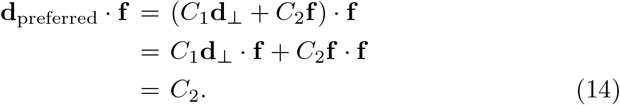

With that, all required parts of SM Equation 8 are determined, letting **d**_actual_ be calculated. Note that if the effective influence parameter *γ* is zero, SM Equation 8 becomes SM Equation 12, meaning **d**_actual_ will equal **d**_preferred_. Likewise, when *γ* is one, its maximum value, **d**_actual_ equals **f**.

In the case that **d**_preferred_ is parallel to **f**, 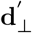 from SM Equation 11 is the zero vector. We assume the norm of the zero vector to be zero. With that set, the calculations will proceed as normal.

Note that there are two possibilities on how to interpret **d**_actual_:

**A:** The cell senses its environment, determines direction of travel along the fibers, and all cell effort goes into that motion.

**B:** The cell attempts to move both along the fibers and along its preferred direction, wasting effort in the process.

These two interpretations lead to either normalizing **d**_actual_ (Option **A**) or letting the cell direction vector lose magnitude by not normalizing **d**_actual_ (Option **B**). For the purposes of this work, we choose Option **B** and do not normalize **d**_actual_.

#### Constitutive relations to determine d_preferred_

To determine **d**_preferred_ in SM Equation 8, we use either the motility model of [40]’s motility model or an adaptation that directly incorporates anisotropy.

##### Model I: Chemotaxis based motility

To determine **d**_preferred_ for chemotactically motivated cells, we use the following equation from [40]:

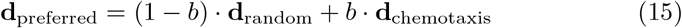

where **d**_random_ is a randomly generated unit vector, **d**_chemotaxis_ is a unit length microenvironment-derived direction, such as the gradient of a cell-required substrate, and the scalar *b*, running from 0 - 1, is the directional bias parameter that sets the balance between random (*b*=0) and completely biased cell motility r, (*b*=1). **d**_preferred_ is the cell-selected motility direction and may be combined with local fiber orientation to produce the final cell motility direction.

##### Model II: ECM and chemotaxis based motility

To couple chemotactic behavior with reading of ECM signals, we reformulated Equation 15 and replace the chemotactic bias with the local ECM anisotropy:

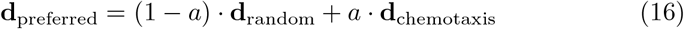

Here, *a* replaces the bias parameter in setting the random versus chemotactic behavior. When combined with the cell-ECM interactions, this model effectively enables a cell to only be chemotactic when in an element of anisotropic ECM.

In either model, **d**_preferred_ is updated stochastically, with an average update interval equal to the cell’s persistence time.

#### 7.2.2 ECM density

We use the following relationship to incorporate the influence of ECM density *ρ* on cell speed:

**Supplementary Figure 2:**
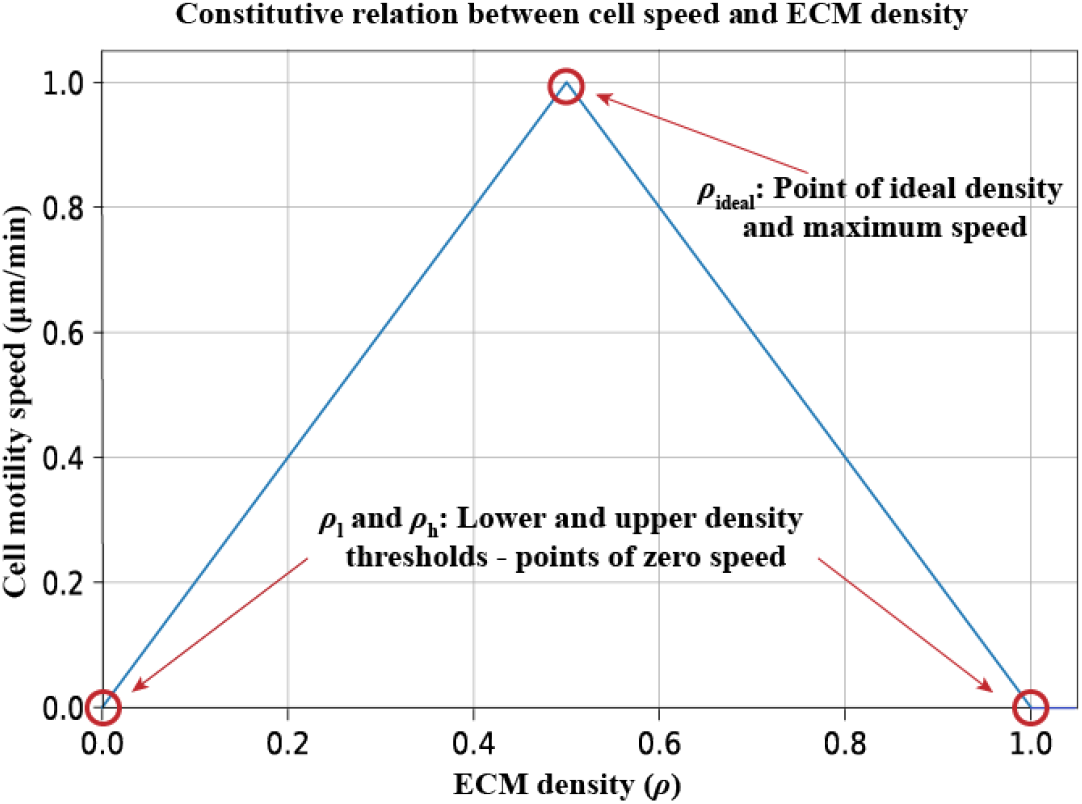
Constitutive relationship between cell speed and density. This is the graphical form of SM Equation 17 with parameters used for the example models.

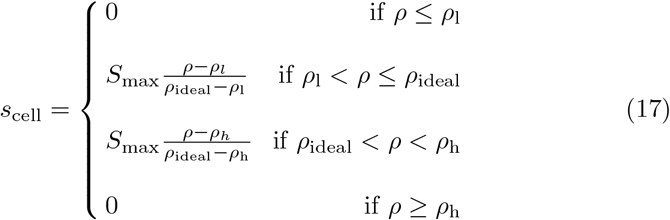

where *ρ* is the local ECM density, *S*_max_ is the maximum cell migration speed, *ρ*_l_ is the ECM threshold density above which cells have non-zero speed, *ρ*_h_ is the ECM threshold density above which cells cannot move, and *ρ*_ideal_ is the ECM density of maximum speed. Supplementary Figure 2 shows the tent-like shape of SM Equation 17 with the parameters used in our example models.

This form simulates a cell that in a fibrous environment cannot move either when density is too low, due to a lack of fibers to grab onto for propulsion, or when density is too high, meaning the ECM is too dense for the cell to pass through. Other polynomials or curves derived from fitting to data could be used in place of this form. We choose it to be a minimum example of a tunable, potentially asymmetric curve with a local maximum.

#### 7.2.3 Combined ECM microstructure and chemical environment influence on cell motility

Combining all these influences together produces the equation for cell velocity due to cell motility (locomotion):

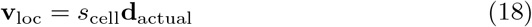

where *s*_cell_ is from SM Equation 17 and **d**_actual_ is from SM Equation 8.

### 7.3 Cell velocity and cell-cell adhesion and repulsion model

In PhysiCell, the agent-based modeling framework in which our examples and ECM model are built, cell velocities and positions update based on several forces - cell-cell forces as well as cell-environmental forces and cell-locomotive force. In this current work, we focused on cell-cell forces (cell-adhesion and repulsion) as well as cell locomotive forces. To discuss this, we follow the explanation from [40], beginning with the cell equation of motion. See [40] and [84] for additional details and discussion.

Given a cell *i* at position **x**_*i*_(*t*), velocity **v**_*i*_(*t*) and a set of neighboring cells 𝒩 (*i*), PhysiCell models cell motion with this equation:

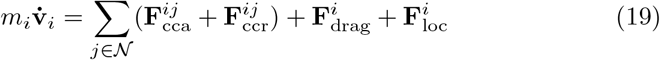

where **F**_cca_ and **F**_ccr_ are the cell-cell adhesive and repulsive (cell resistance to deformation) forces, **F**_drag_ represents dissipative, drag-like forces such as viscous drag, and **F**_loc_ is cell generated, or motile, force. Note that as appropriate, additional force terms can be added (see [40] and [84]).

Examining this term by term, drag is modeled as follows [40, 84, 85]:

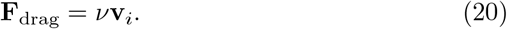

Here, *ν* is a drag coefficient. This differs from the formulation in [40], which also included a drag term for ECM interactions. Since we model ECM interactions explicitly in the update for cell locomotive force, we do not it include it here. The formulation makes the inertialess assumption 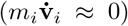 [40, 84, 86, 87] which assumes that any changes to forces on the cell rapidly equilibrate. Making these substitutions in Equation 19 and solving for **v**_*i*_, we get:

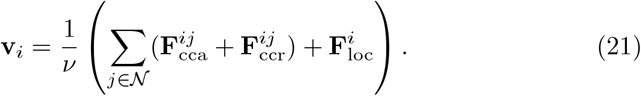

As noted in [84], this assumption yields the interpretation of each term as the terminal velocity of the cell, given that only the force of interest and drag are acting on the cell. The locomotive force and its contribution to cell velocity is covered in the previous section (7.2). The forces **F**_cca_ and **F**_ccr_ are then modeled using interaction potentials that are functions of maximum adhesion distance, cell geometry (radius), adhesion and repulsion parameters, and distance to other cells. The adhesion function is:

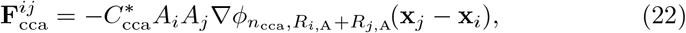

where 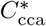 is the cell-cell adhesion parameter, *A*_*i*_ and *A*_*j*_ are the cell-cell relative adhesion parameters (bounded inclusively between 0 and 1), *φ* is the potential function, *n*_cca_ a parameter setting the shape of the potential function, and *R*_*i*,A_ and *R*_*j*,A_ are the maximum adhesion distances of cells *i* and *j* respectively. Note that this gives 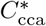 units of force. To define the mechanics parameter in PhysiCell, we divide through by *ν* giving:

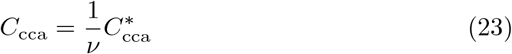

with *C*_cca_ taking on units of speed, as expected for SM Equation 21.

This is the repulsion function:

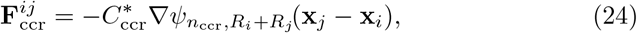

where 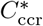 is the cell-cell repulsion parameter, *n*_ccr_ is a parameter setting the shape of the potential function, and *R*_*i*_ and *R*_*j*_ are the cell radii of the two interacting cells *i* and *j*. As above with 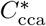, we divide 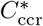 by *ν*:

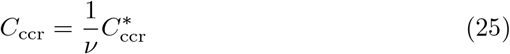

giving *C*_ccr_ units of speed.

Finally, PhysiCell uses the following potential functions for adhesion and repulsion respectively:

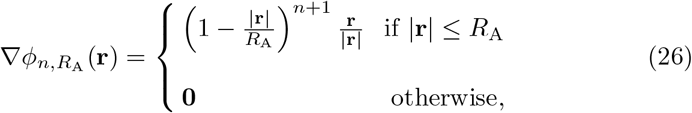

and:

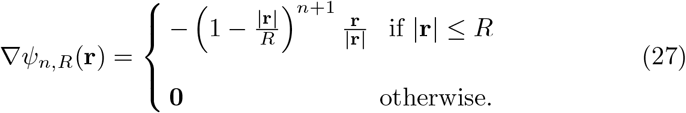

Note that to compute the velocity updates, only the gradients of the potentials are required. **r** is the vector displacement between the interacting cells’ centers, *R*_A_ is the sum of *i* and *j*’s interaction distances, and *R* is the sum of the two cells’ diameters.

### 7.4 PhysiCell rules

PhysiCell rules is a new extension to PhysiCell [62]. It uses a grammar to encode a mathematical representation of cell-based rules or hypotheses for an arbitrary number of signals (*e*.*g*. - contact with other cells, a substrate concentration, ECM density) increasing or decreasing a cell behavior (*e*.*g*. – cell speed, cell-cell adhesion, ECM production rate). The base form is as follows:

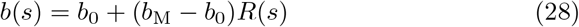

where *b*_0_ is the base level of a rate or other parameter modifiable at the cell level, *b*_M_ is the maximum quantity expected, and *R*(*s*) is the functional relationship between the behavior *b* and signal *s*. In our examples, we use the default functional form for *R*(*s*) - Hill functions. Each Hill function requires a half-maximum saturation value and Hill exponent. As more than one signal may influence a behavior and the influence may be excitatory or inhibitory, Johnson et al. generalize the response of behavior *b* to a vector of **u** and **d** up and down signals respectively. These are used to produce the general form of Equation 28:

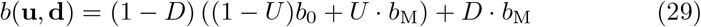

where *D* and *U* are the total up and down responses to vectors of signals **u** and **d**.

### 7.5 Model parameters

We performed a joint parameter sweep to determine the cell-cell adhesion and cell speeds using a preliminary version of the collective invasion model. Those initial values were modified for the additional example models. Similarly, the fiber realignment and reorientation rates were initially determined through single variable parameter sweep of preliminary versions of the collective invasion model. Parameters related to oxygen fields are from previous literature. Additional parameter values were selected to enable emergent behavior to occur at a time scale on the order of simulated days.

#### 7.5.1 Invasive cellular front parameter details

See Tables 4 to 6 for the invasive cellular front parameters. Note there is no chemical environment or rules for this model.

**Table 4:** Cell-level parameters specific to the invasive cellular front model.

| Parameter | Value |
| --- | --- |
| Adhesion strength | 0.4 $\mu\text{m}/\text{min}$ |
| Repulsion strength | 25.0 $\mu\text{m}/\text{min}$ |
| Base cell speed | 1.25 $\mu\text{m}/\text{min}$ |
| $\rho_{\text{ideal}}$ | 0.5 |
| $\rho_{\text{h}}$ | 1.0 |
| $\rho_{\text{l}}$ | 0.0 |
| ECM sensitivity | 1.0 |
| Rate of proliferation | 0 $\text{min}^{-1}$ |
| Rate of death | 0 $\text{min}^{-1}$ |
| Chemotactic bias | 0.0 |
| Persistence time | 10 min |
| Fiber realignment rate (rate of change in anisotropy) | 0.0 $\text{min}^{-1}$ |
| Fiber reorientation rate | 0.0 $\text{min}^{-1}$ |
| Density modification rate | 0.0 $\text{min}^{-1}$ |

**Table 5:** ECM initial conditions for invasive cellular front model.

| Parameter | Value |
| --- | --- |
| Initial anisotropy | 1.0 |
| Initial density | 0.5 |
| Initial orientation | random, horizontal, vertical, or mixed horizontal and vertical |

**Table 6:** Tissue level parameters for invasive cellular front model.

| Parameter | Value |
| --- | --- |
| Computational domain size | 600 $\mu\text{m}$ by 1000 $\mu\text{m}$ |
| Number of cells added per cell addition step | 30 cells |
| Cell addition step size | 180 min |

#### 7.5.2 Fibrosis parameter details

See Tables 7 to 10 for the fibrosis model parameters. Rules are below.

**Table 7:** Cell-level parameters specific to the fibrosis model.

| Parameter | Value |
| --- | --- |
| Adhesion strength | Fibroblast and macrophage - 5.0 $\mu\text{m}/\text{min}$, Dead cell - 10.0 $\mu\text{m}/\text{min}$ |
| Repulsion strength | 25.0 $\mu\text{m}/\text{min}$ |
| Base cell speed | Fibroblast and macrophage - 0.5 $\mu\text{m}/\text{min}$, Dead cell - 0.0 $\mu\text{m}/\text{min}$ |
| $\rho_{\text{ideal}}$ | All - 0.5 |
| $\rho_{\text{h}}$ | All - 1.0 |
| $\rho_{\text{l}}$ | All - 0.0 |
| ECM sensitivity | All - 0.0 |
| Debris secretion rate | Dead cells - 1.0 $\text{min}^{-1}$ |
| Debris target | Dead cells - 10 |
| Debris uptake | Macrophages - 1.0 $\text{min}^{-1}$ |
| Inflammatory signal secretion rate | Macrophages - see Fibrosis rules (Section 7.5.2) |
| Inflammatory signal target | Macrophages - 1.0 |
| Rate of proliferation | All - 0 $\text{min}^{-1}$ |
| Rate of death | All - 0 $\text{min}^{-1}$ (dead cells begin the simulation already in a death process) |
| Transition rate from death state to removed from simulation | Dead cells - 0.001938 $\text{min}^{-1}$ |
| Chemotactic bias | Fibroblasts and macrophages - 0.5 |
| Persistence time | 10 min |
| Fiber realignment rate (rate of change in anisotropy) | Fibroblast - 0.004 $\text{min}^{-1}$ |
| Fiber reorientation rate | Fibroblast - 4 $\text{min}^{-1}$ |
| $\rho_{\text{target}}$ | Macrophages - 0.5, Fibroblast - 1.0 |
| Density modification rate | Macrophages - 0.0 $\text{min}^{-1}$, Fibroblast - see Fibrosis rules (Section 7.5.2) |

**Table 8:**
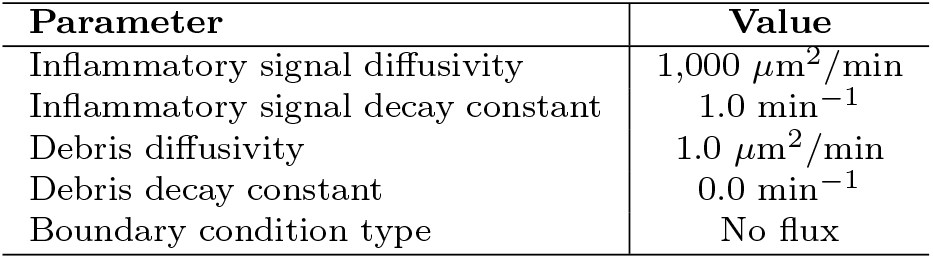
Biotransport and chemical microenvironment parameters for fibrosis model.

**Table 9:**
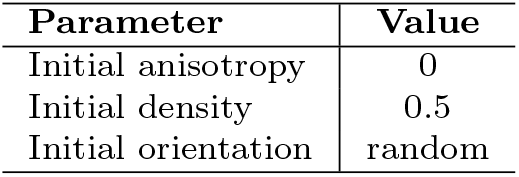
ECM initial conditions for fibrosis model.

| Parameter | Value |
| --- | --- |
| Initial anisotropy | 0 |
| Initial density | 0.5 |
| Initial orientation | random |

**Table 10:** Tissue level parameters for fibrosis model.

| Parameter | Value |
| --- | --- |
| Initial region of distressed/dead cell radius | 175 $\mu\text{m}$ |
| Computational domain size | 1600 $\mu\text{m}$ by 1600 $\mu\text{m}$ |
| Pre-mechanics microenvironment conditioning duration | 10 min |

##### Fibrosis cell rule details

In fibroblast cells:

- Contact with macrophage increases `custom:ECM_production_rate` (ECM density production) rate from 0.0001 min^−1^ towards 0.001 min^−1^ with a Hill response, with half-maximum 0.1 contacts and Hill power 10.

In dead cells:

- None

In macrophage cells:

- Contact with dead cell increases `inflammatory_signal` secretion from 0 towards 10 min^−1^ with a Hill response, with half-maximum 0.1 contacts and Hill power 10.
- Volume decreases `phagocytose dead cell` from 0.0005 min^−1^ towards 0.0001 min^−1^ with a Hill response, with half-maximum 2494 *µ*m^3^ and Hill power 10.

#### 7.5.3 Basement membrane degradation parameter details

See Tables 11 to 14 for the invasive carcinoma parameters. Rules are below.

**Table 11:**
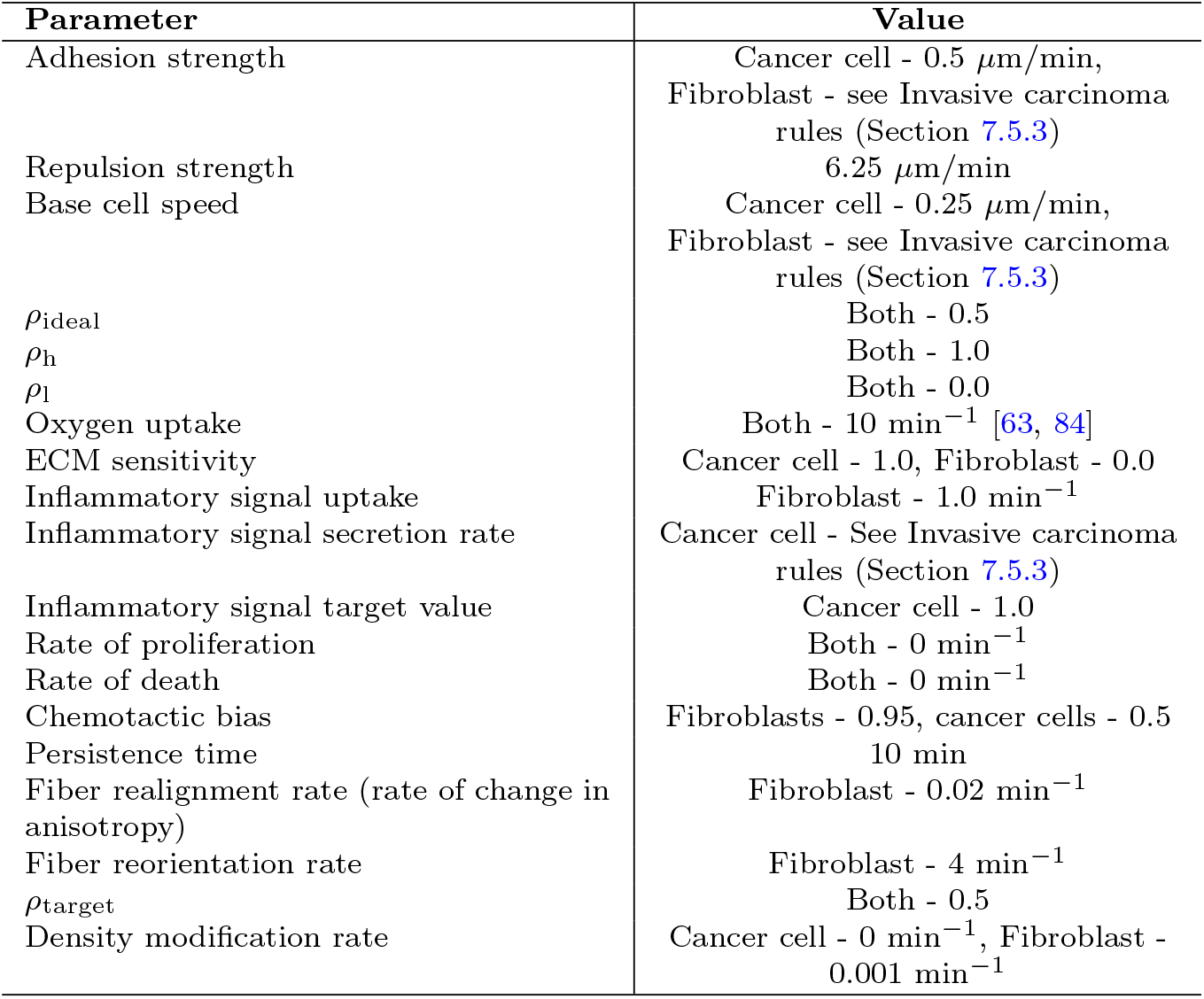
Cell-level parameters specific to the basement membrane degradation model.

| Parameter | Value |
| --- | --- |
| Adhesion strength | Cancer cell - 0.5 $\mu\text{m}/\text{min}$,<br>Fibroblast - see Invasive carcinoma rules (Section 7.5.3) |
| Repulsion strength | 6.25 $\mu\text{m}/\text{min}$ |
| Base cell speed | Cancer cell - 0.25 $\mu\text{m}/\text{min}$,<br>Fibroblast - see Invasive carcinoma rules (Section 7.5.3) |
| $\rho_{\text{ideal}}$ | Both - 0.5 |
| $\rho_{\text{h}}$ | Both - 1.0 |
| $\rho_{\text{l}}$ | Both - 0.0 |
| Oxygen uptake | Both - 10 $\text{min}^{-1}$ [63, 84] |
| ECM sensitivity | Cancer cell - 1.0, Fibroblast - 0.0 |
| Inflammatory signal uptake | Fibroblast - 1.0 $\text{min}^{-1}$ |
| Inflammatory signal secretion rate | Cancer cell - See Invasive carcinoma rules (Section 7.5.3) |
| Inflammatory signal target value | Cancer cell - 1.0 |
| Rate of proliferation | Both - 0 $\text{min}^{-1}$ |
| Rate of death | Both - 0 $\text{min}^{-1}$ |
| Chemotactic bias | Fibroblasts - 0.95, cancer cells - 0.5 |
| Persistence time | 10 min |
| Fiber realignment rate (rate of change in anisotropy) | Fibroblast - 0.02 $\text{min}^{-1}$ |
| Fiber reorientation rate | Fibroblast - 4 $\text{min}^{-1}$ |
| $\rho_{\text{target}}$ | Both - 0.5 |
| Density modification rate | Cancer cell - 0 $\text{min}^{-1}$, Fibroblast - 0.001 $\text{min}^{-1}$ |

**Table 12:**
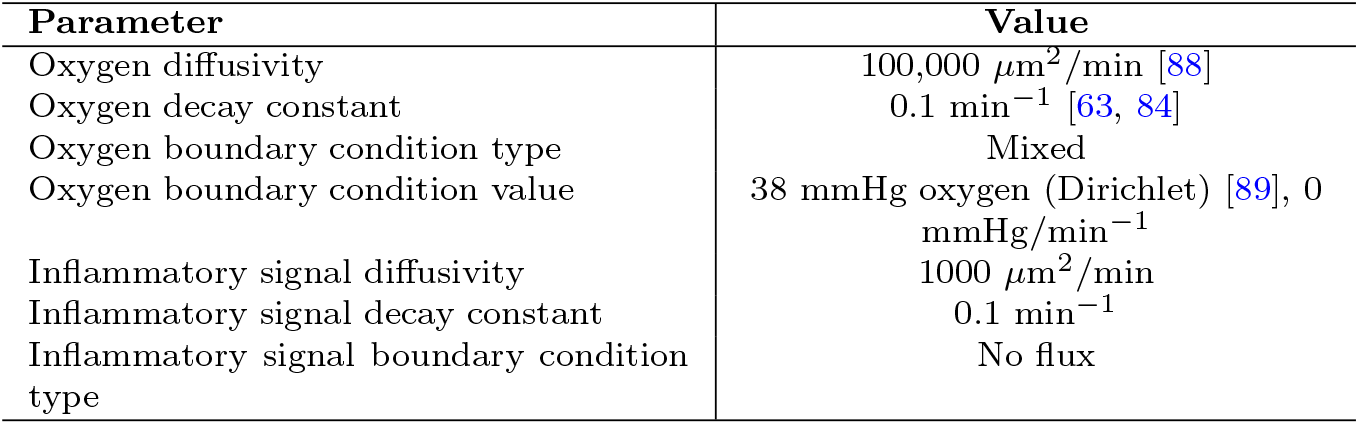
Biotransport and chemical microenvironment parameters for basement membrane degradation model.

| Parameter | Value |
| --- | --- |
| Oxygen diffusivity | 100,000 $\mu\text{m}^2/\text{min}$ [88] |
| Oxygen decay constant | 0.1 $\text{min}^{-1}$ [63, 84] |
| Oxygen boundary condition type | Mixed |
| Oxygen boundary condition value | 38 mmHg oxygen (Dirichlet) [89], 0 $\text{mmHg}/\text{min}^{-1}$ |
| Inflammatory signal diffusivity | 1000 $\mu\text{m}^2/\text{min}$ |
| Inflammatory signal decay constant | 0.1 $\text{min}^{-1}$ |
| Inflammatory signal boundary condition type | No flux |

**Table 13:**
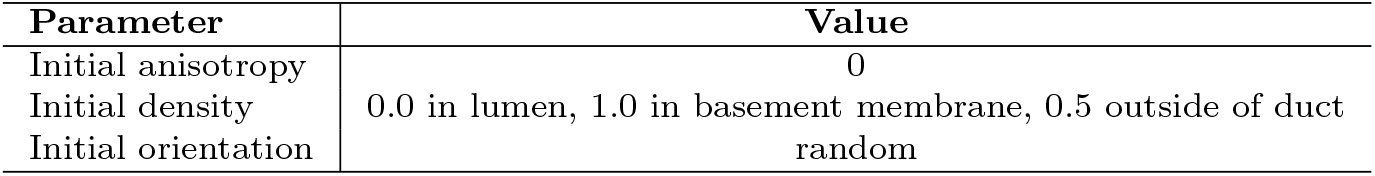
ECM initial conditions for the basement membrane degradation model.

| Parameter | Value |
| --- | --- |
| Initial anisotropy | 0 |
| Initial density | 0.0 in lumen, 1.0 in basement membrane, 0.5 outside of duct |
| Initial orientation | random |

**Table 14:** Tissue level parameters for basement membrane degradation model.

| Parameter | Value |
| --- | --- |
| Initial tumor radius | 175 $\mu\text{m}$ |
| Computational domain size | 1600 $\mu\text{m}$ by 1600 $\mu\text{m}$ |
| Pre-mechanics microenvironment conditioning duration | 10 min |

##### Basement membrane degradation rule details

In fibroblast cells:

- Contact with cancer cell decreases `custom:rules_based_speed_multiplier`, a dimensionless multiplier that reduces or increases base cell speed, from 1 towards 0.5 with a Hill response, with half-maximum 0.1 contacts and Hill power 4.

In cancer cells:

- contact with fibroblast decreases `inflammatory_signal` secretion from 50 min^−1^ towards 1 with a Hill response, with half-maximum 0.5 contacts and Hill power 4.
- Contact with fibroblast decreases `adhesive affinity to cancer cell` from 1 towards 0.25 with a Hill response, with half-max 0.1 and Hill power 4.
- Contact with fibroblast decreases `cell-cell adhesion` from 0.5 *µ*m/min towards 0.25 *µ*m/min with a Hill response, with half-max 0.1 contacts and Hill power 4.

#### 7.5.4 Collective migration parameter details

See Tables 15 to 18 for collective migration parameters. Note there are no cell rules for this model.

**Table 15:** Cell-level parameters specific to the collective migration model. Unless otherwise stated, parameters are at default values.

| Parameter | Value | Range |
| --- | --- | --- |
| Adhesion strength | varies | 0, 10 $\mu\text{m}/\text{min}$ |
| Repulsion strength | 25.0 $\mu\text{m}/\text{min}$ | - |
| Base cell speed | varies | 0.1, 0.5, 0.8 $\mu\text{m}/\text{min}$ |
| $\rho_{\text{ideal}}$ | 0.5 | - |
| $\rho_{\text{h}}$ | 1.0 | - |
| $\rho_{\text{l}}$ | 0.0 | - |
| ECM sensitivity | varies | 0.0 (no reading), 1.0 (all other simulations), 0 for leaders (all simulations) |
| Oxygen uptake | 10 $\text{min}^{-1}$ [63, 84] | - |
| Rate of proliferation | 0 $\text{min}^{-1}$ | - |
| Rate of death | 0 $\text{min}^{-1}$ | - |
| Chemotactic bias | Leaders - 0.95, followers - dynamic | - |
| Persistence time | 10 min | - |
| Fiber realignment rate (rate of change in anisotropy) | varies | 0.0 (no writing), 0.001 (loss of collective migration scenario), 0.004 $\text{min}^{-1}$, 0 for followers (all simulations) |
| Fiber reorientation rate | varies | 0.0 (no writing), 1 (loss of collective migration scenario), 4 $\text{min}^{-1}$, 0 for followers (all simulations) |
| $\rho_{\text{target}}$ | 0.5 | - |
| Density modification rate | 0.0 | - |

**Table 16:** Biotransport and chemical microenvironment parameters for collective migration model.

| Parameter | Value |
| --- | --- |
| Oxygen diffusivity | 100,000 $\mu\text{m}^2/\text{min}$ [88] |
| Oxygen decay constant | 0.1 $\text{min}^{-1}$ [63, 84] |
| Boundary condition type | Dirichlet |
| Boundary condition value | 38 mmHg oxygen [89] |

**Table 17:**
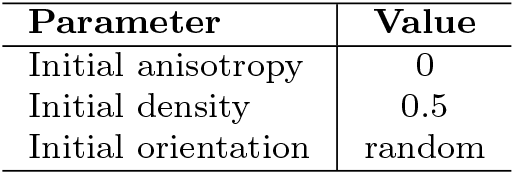
ECM initial conditions for all collective migration simulations.

**Table 18:** Tissue level parameters for collective migration model.

| Parameter | Value |
| --- | --- |
| Initial organoid radius | 175 $\mu\text{m}$ |
| Computational domain size | 1600 $\mu\text{m}$ by 1600 $\mu\text{m}$ |
| Pre-mechanics microenvironment conditioning duration | 10 min |
| Cell fraction of leaders | 0.05 |

**Table 19:** Summary statistics of stochastic replicates of the invasive front simulation. We present the mean and standard deviation of the bin centers (y-coordinate) containing the 95th percentile of the cell count across all 21 replicates per scenario. For reference, the y-coordinates in this simulation runs from -500 *µ*m to 500 *µ*m.

| Scenario | Mean ($\mu\text{m}$) | Standard Deviation ($\mu\text{m}$) |
| --- | --- | --- |
| Random | -124.4 | 12.5 |
| Parallel | -162.0 | 0.0 |
| Perpendicular | 1.8 | 21.2 |

### 7.6 Links to supplementary videos

- Writing to ECM (local microstructure remodeling) - Cell-based ECM reorientation (Figure 2a)
- Reading ECM (contact guidance) - Circular ECM with no gradient (Figure 2b)
- Reading ECM (contact guidance) - Circular ECM with chemical gradient (Figure 2c)
- Invasive cell front - randomly oriented ECM (Figure 3)
- Invasive cell front - parallel-oriented ECM (Figure 3)
- Invasive cell front - perpendicular-oriented ECM (Figure 3)
- Invasive cell front - mixed ECM (Figure 3)
- Wound healing and fibrosis (Figure 4)
- Basement membrane degradation and transition of *in situ* carcinoma to invasion (Figure 5)
- Writing signals to ECM only (no reading/contact guidance) (Figure 6a)
- Reading signals from ECM only (no writing/microscale remodeling) (Figure 6b)
- Local microstructure remodeling (reading) and writing (contact guidance)
- leads to stigmergy (Figure 6c)
- Leader-follower - instant remodeling - higher cell speed (Figure 7a)
- Leader-follower - instant remodeling - medium cell speed (Figures 7b, 8a, SM4a)
- Leader-follower - instant remodeling - lower cell speed (Figure 7c)
- Collective migration - Non-instant signal writing (Figure 8b)
- Reading ECM (contact guidance) - split ECM pattern with chemical gradient (SM Figure 3)
- Collective migration - Loss of collective migration due to decreased remodeling rates (SM Figure 4b)

### 7.7 Supplementary Figures and invasive front results details

**Supplementary Figure 3:**
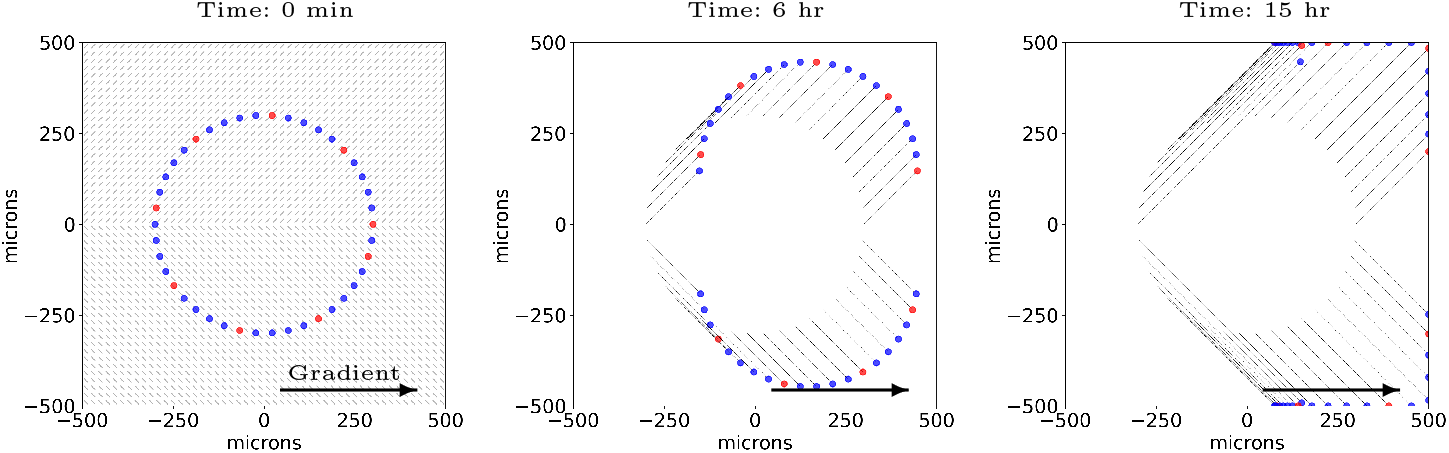
Highlight of individual aspects of the ECM-cell interactions: ECM orientation and cell chemotaxis. A combination of cues directing cell motility: ECM orientation (45^*°*^ in the top of the domain and -45^*°*^ in the bottom) and chemical gradient (to the right). The small black arrows trailing the cells show the positional history of each cell. The cell position is marked every six simulated minutes. Red and blue cells are identical; the coloring was added only for visual contrast. This is available as a video. See link the to video here.

**Supplementary Figure 4:**
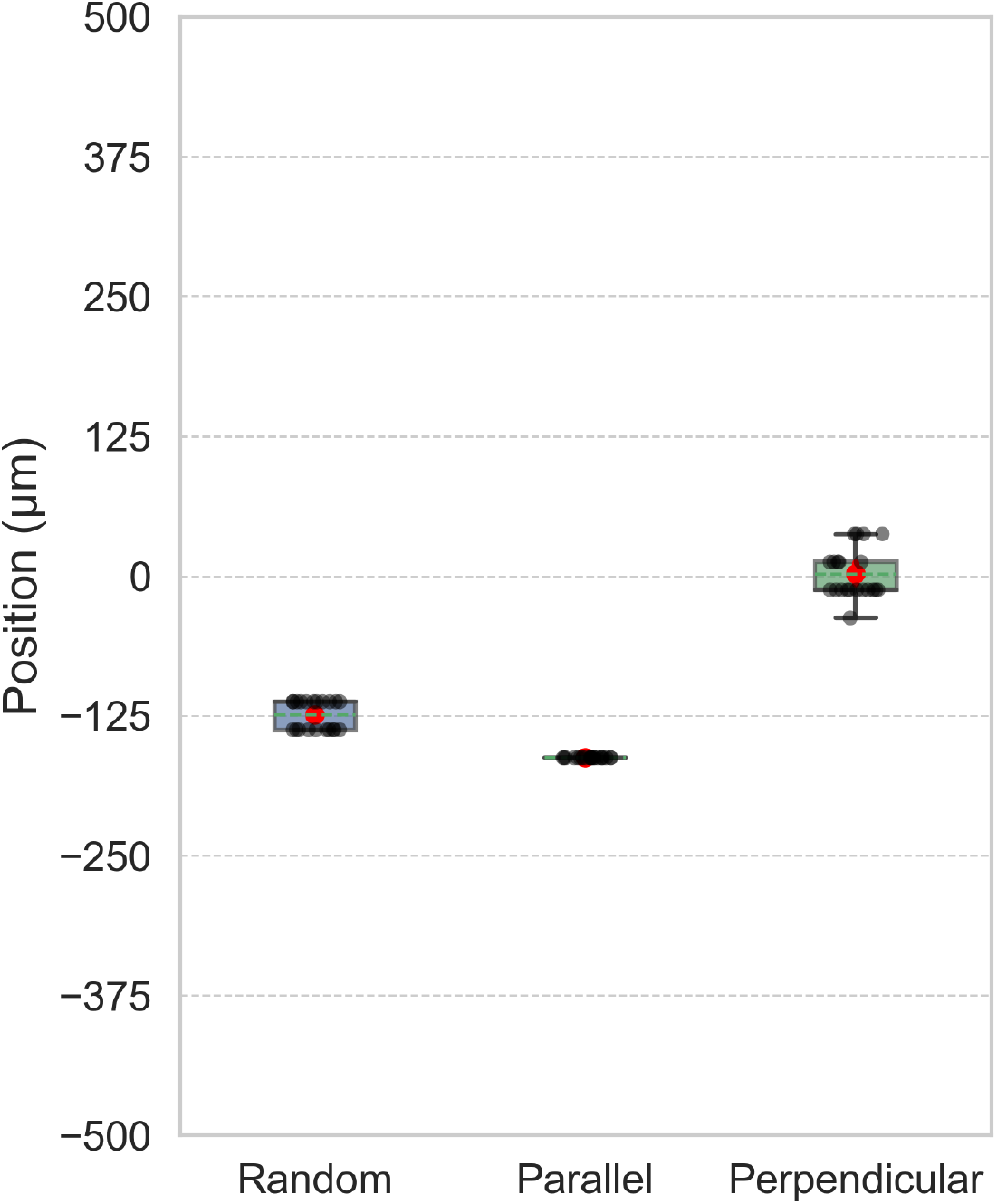
Extent of invasive front in invasive front scenarios: Jitter and box and whisker plots of results across the random, parallel, and perpendicular ECM orientation scenarios. Each circle shows the y-position of the histogram bin containing the 95th percentile of cell count at five simulated days. The red circle marks each distributions’ mean. Box and whiskers mark distribution quartiles.

**Supplementary Figure 5:**
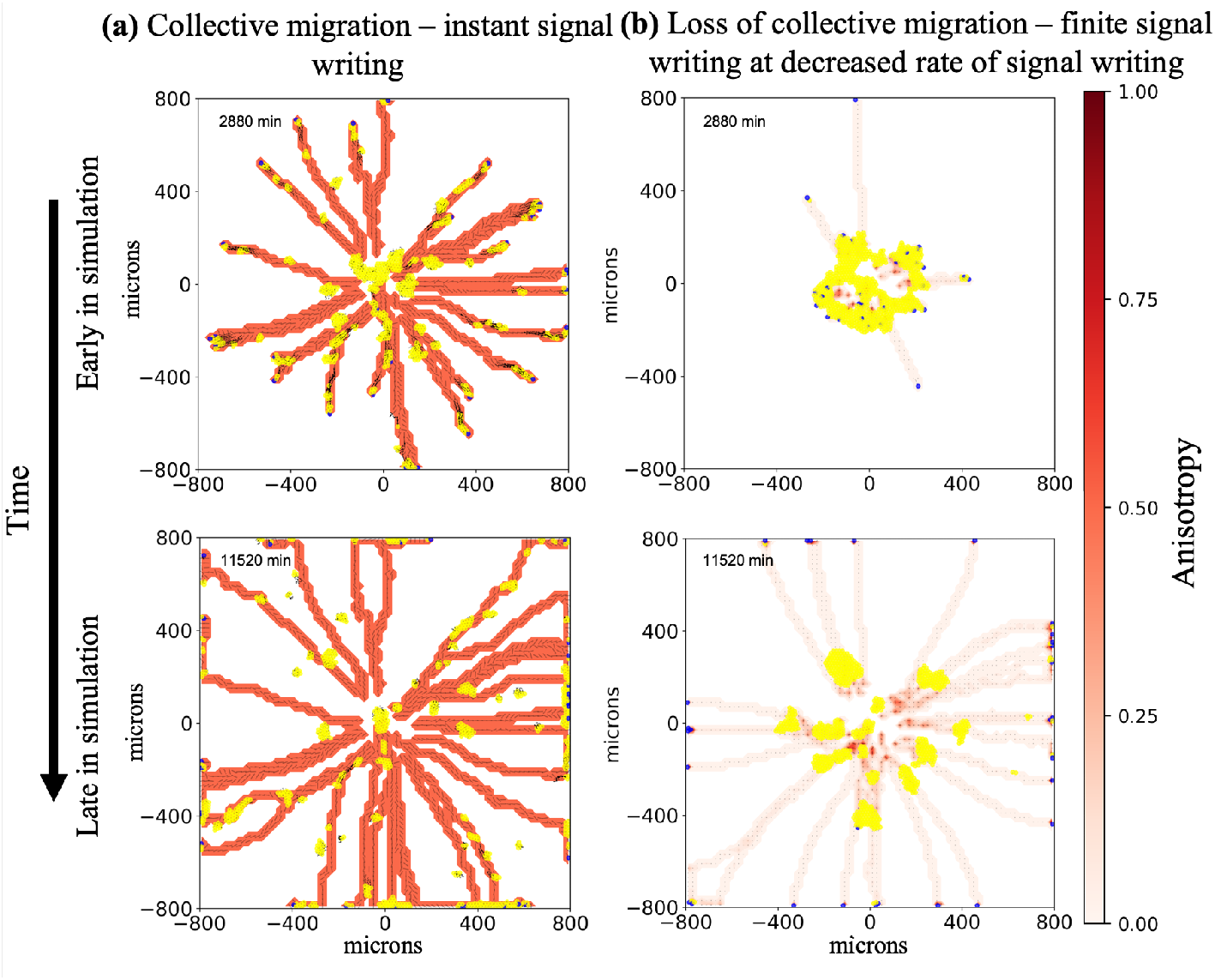
Collective migration requires sufficiently fast remodeling parameters: Collective cell migration is observed in **(a)**, which has instantaneous ECM remodeling, whereas **(b)**, which has relaxed the instantaneous signal writing and slower rates of remodeling than those in Figure 8b, lacks collective migration. Fiber realignment rate: 0.001 min^−1^, fiber reorientation rate: 1 min^−1^. Videos are available. See the links here.

